# Therapeutic Potential of iPSC-Derived TH^⁺^/FOXA2^⁺^ Neuronal Extracellular Vesicles in Parkinson’s Disease Revealed by Organoid and Rodent Models

**DOI:** 10.64898/2026.09.22.753408

**Authors:** Zhen Qi, Xiaowen Li, Xiao Jin, Chen Shen, Lan Yang, Yandan Liu, Mingqian Huang, Lina Xing, Yan Chen, Zongjun Liu

## Abstract

**Background:** Parkinson’s disease (PD) is characterized by the progressive loss of midbrain dopaminergic (DA) neurons and aberrant α-synuclein (α-syn) aggregation, yet disease-modifying therapies that simultaneously retard neurodegeneration and foster regeneration remain lacking.

**Methods:** In this study, we isolated extracellular vesicles (TH⁺/FOXA2⁺ cell-derived EVs) from TH⁺/FOXA2⁺ differentiated neural cells, a distinct midbrain floor-plate-derived neural cell population that co-expresses the floor-plate transcription factor FOXA2 together with TH and should therefore not be equated with conventional mature midbrain dopaminergic neurons, and systematically evaluated their therapeutic potential across three complementary models: A53T transgenic mice, 6-OHDA-lesioned-rats, and 6-OHDA-treated human midbrain organoids. Integrative transcriptomic and single-cell-sequencing analyses, together with toxicological evaluations in rodents and non-human primates, were performed to investigate the underlying mechanisms and safety profile. These TH⁺/FOXA2⁺ cells were generated using a fully suspension-based, matrix-free, and chemically defined differentiation system devoid of N2/B27 supplements and fetal bovine serum (FBS). This approach offers a simple, controllable, low-cost, and scalable platform with high batch-to-batch consistency for EV manufacturing. Throughout this study, the term TH⁺/FOXA2⁺ cells refers to this distinct TH⁺/FOXA2⁺ cell population and should not be interpreted as conventional terminally differentiated dopaminergic neurons.

**Results:** Intranasal administration of TH⁺/FOXA2⁺ cell-derived EVs in A53T mice and 6-OHDA-lesioned rats significantly ameliorated motor deficits, increased nigral tyrosine hydroxylase (TH)-positive-neuron counts, reduced α-syn-aggregation, and suppressed gliosis. In human midbrain organoids, TH⁺/FOXA2⁺ cell-derived EVs preserved DA neuronal morphology and reduced GFAP and α-syn expression. Notably, when EVs were administered simultaneously with 6-OHDA modeling rather than after modeling was completed, the protective effect was stronger. Multi-omics-analysis revealed that TH⁺/FOXA2⁺ cell-derived EVs reversed pathological signatures of interferon-responsive-microglia and oxidative stress-adapted astrocytes, while restoring WNT and non-canonical-WNT mediated intercellular communications. Safety evaluations showed no discernible organ toxicity.

**Conclusions:** TH⁺/FOXA2⁺ cell-derived EVs exert neuroprotective effects across multiple PD models by modulating specific glial subpopulations and reinstating development-associated signaling pathways. Together with a fully suspension, matrix-free and N2/B27-free differentiation process that enables low-cost, scalable and highly reproducible manufacturing, TH⁺/FOXA2⁺ cell-derived EVs represent a promising therapeutic candidate for Parkinson’s disease.-

## 1. Introduction

Parkinson’s disease (PD) is the second most common neurodegenerative disorder worldwide. Its core pathological feature is the progressive loss of dopaminergic neurons in the substantia nigra pars compacta of the midbrain, leading to severe dopamine deficiency in the striatum[1, 2]. At the molecular pathological level, abnormal aggregation of α-synuclein (α-syn) represents one of the most critical hallmarks of the disease[3]. α-Synuclein is a protein predominantly expressed at presynaptic terminals, where it participates in the regulation of neurotransmitter release and synaptic plasticity under physiological conditions. However, under the pathological conditions of Parkinson’s disease, factors such as genetic mutations, oxidative stress, or dysfunction of the protein clearance system cause soluble α-syn monomers to misfold into oligomers and insoluble fibrils, ultimately forming the characteristic eosinophilic intracytoplasmic inclusions known as Lewy bodies[4]. This aggregation process not only directly exerts protein toxicity but, more importantly, confers upon misfolded α-syn a prion-like “seed” property that enables cell-to-cell propagation, inducing conformational changes in normal α-syn and thereby driving the retrograde spread of pathology from peripheral/enteric sites to the substantia nigra and cerebral cortex[5]. Concomitantly, the expression of its terminal marker enzyme, tyrosine hydroxylase (TH), is markedly reduced in the striatum, resulting in impaired dopamine synthesis[6]. Similarly, oxidative stress plays an equally critical role in sporadic Parkinson’s disease as well as in toxin-induced-models. 6-Hydroxydopamine-(6-OHDA-), a classical neurotoxin, is specifically taken up by dopaminergic neurons via the dopamine transporter. Through auto-oxidation-, it generates reactive oxygen species and quinones, directly inhibiting mitochondrial complex I and inducing lipid peroxidation, ultimately leading to neuronal apoptosis[7, 8]. Consequently, the 6-OHDA-lesioned--rat has become a classic tool model for recapitulating oxidative stress-induced-injury in sporadic PD[9–12]. Therefore, targeting abnormal α-syn-aggregation or alleviating oxidative stress-induced-neuronal damage is considered a key strategy to reverse the course of PD and promote neural regeneration. Given the etiological heterogeneity of PD, cross-validation-using both genetic and oxidative stress models enables a more comprehensive assessment of the generalizability of potential therapeutic interventions.

In recent years, stem cell- and stem cell-derived-therapies have shown great promise in regenerative medicine[13]. Among these, extracellular vesicles (EVs), as key mediators of intercellular communication, have been recognized as ideal vehicles for “cell-free-therapy” due to their low immunogenicity, ability to carry bioactive molecules (e.g., miRNAs, proteins), and capacity to readily cross biological barriers[14]. Notably, EVs derived from different cell sources exhibit distinct homing capabilities and cargo profiles. Studies have demonstrated that brain organoid-derived-EVs are superior to mesenchymal stem cell-derived-EVs in promoting the differentiation of human induced pluripotent stem cells (iPSCs) into dopaminergic neurons. This advantage is attributed to the higher abundance of neurotrophic factors and glial cell line-derived-neurotrophic factor (GDNF) in brain organoid-derived-EVs, which facilitates the differentiation of iPSCs into DA neurons in a LIM homeobox transcription factor 1 alpha (LMX1A)--dependent manner[15]. Therefore, it is theoretically plausible that EVs originating from the dopaminergic neuron microenvironment may preferentially target the nigrostriatal pathway and carry key trophic factors essential for the survival of dopaminergic neurons. However, efficient and non-invasiv-e delivery of EVs to specific brain regions remains a major challenge for clinical translation. Intranasal administration, a non-invasiv-e method that bypasses the blood–brain barrier and delivers drugs directly to the central nervous system via the olfactory and trigeminal pathways, has been widely used in the treatment of neurological disorders, offering new strategies for brain-targete-d therapy[16–18]. Against this background, the present study employed three complementary models – A53T mice, 6-OHDA-lesione--d rats, and 6-OHDA-treate--d midbrain organoids – to evaluate the neuroprotective and neuroregenerative effects of TH⁺/FOXA2⁺ cell-derived EVs on dopaminergic neurons. This integrated system, spanning *in vivo* to *in vitro*, rodent to human cells, and genetic to toxicological models, enables rigorous validation of whether the effects of EVs are consistent across models. By assessing behavioral improvements, the number of TH-positiv-e neurons, α-synuclei-n aggregation, and downstream synaptic protein expression, we aim to explore the underlying mechanisms. This study is expected to provide a safe, convenient, and efficient precision intervention strategy for the clinical treatment of PD.

## 2. Materials and Methods

### 2.1 Preparation of TH⁺/FOXA2⁺ cell-derived EVs

To obtain the target cells, we established a novel suspension differentiation method (Chinese Invention Patent, Application No. CN2025108899748). The detailed protocol is as follows: Human induced pluripotent stem cells (iPSCs) were maintained on Laminin-521-(Novoprotein)-coated-culture plates in mTeSR™ Plus medium. On day 0 of differentiation, iPSCs were dissociated with Gentle Cell Dissociation Reagent (GCDR, STEMCELL Technologies) at 37 °C for 5–10 minutes to obtain a single-cell-suspension. The cell suspension was supplemented with 10μM ROCK inhibitor Y-27632-(MCE), and cells were then seeded at a density of 5,000–10,000 cells per well into ultra-low-attachment 96-well-plates (Corning) to allow spontaneous aggregation into embryoid bodies (EBs). The seeding volume was 100μL per well, and the medium consisted of mTeSR™ Plus containing 10 μM Y-27632-. After static culture of the 96-well-plate in a 37°C, 5% CO₂ incubator for 24 hours, 100μL of mTeSR™ Plus medium was added to each well. On day 2, the medium was replaced with neural induction medium consisting of 1% GlutaMAX™ (Thermo Fisher), 1% MEM non-essential amino acids (Thermo Fisher), 1% sodium pyruvate, 0.1 mM ascorbic acid (Sigma Aldrich), 0.5g/L albumin (Rongsheng), and 5mg/L insulin (Procell). Dual SMAD inhibitors were added simultaneously: 10μM SB431542 (TGF-β inhibitor, Tocris), 100nM LDN193189 (BMP inhibitor, STEMCELL Technologies), and 10 μ M Purmorphamine (Smoothened agonist, MCE) for 3 days of induction. To efficiently induce midbrain cell fate differentiation, from day 3 onward, SB431542 and LDN193189 were removed, and 3 μM CHIR99021 (GSK-3 inhibitor, MCE) was added. The neural induction medium containing the above-mentioned small molecules and cytokines was replaced every two days for a total of 6 days. From day 9 onward, 3μM CHIR99021 and 10 μM Purmorphamine were maintained until day 15, with the additional supplementation of 1 μM Forskolin to support dopaminergic neuron specification. From day 15 onwards, only 1 μM Forskolin was retained, and 0.5μM retinoic acid (Sigma Aldrich) was added to promote neuronal maturation. Around day 25 of differentiation, cells exhibited a TH⁺/FOXA2⁺ dual-positive-phenotype, and their midbrain dopaminergic neuron identity was validated by immunofluorescence staining or flow cytometry. Importantly, apart from the maintenance of undifferentiated iPSCs before the onset of differentiation, the entire differentiation process—from EB formation on day 0 to the harvest of TH⁺/FOXA2⁺ cells—was carried out exclusively in suspension in ultra-low attachment vessels, without any Matrigel or other extracellular matrix coating and without feeder cells. In addition, the differentiation medium used throughout was chemically defined and completely free of N2 and B27 supplements. By eliminating costly matrix proteins and proprietary supplements, and by reducing manual handling steps, this system constitutes a simple and controllable suspension culture platform featuring low reagent cost, ready scalability and high batch-to-batch consistency, thereby supporting reproducible large-scale production of TH⁺/FOXA2⁺ cell-derived EVs.

During the D30–D60 time window, cell culture supernatants were collected every two days. The supernatants were first clarified by filtration through a 0.45μm pore-size-filter to effectively remove residual cell debris and large-sized-particles. The clarified supernatants were then concentrated using a tangential flow filtration (TFF) system with a molecular weight cut-off of 100kDa at 4 °C. During the concentration process, the feed flow rate and transmembrane pressure were controlled to achieve a final concentrated volume of 1/50 to 1/100 of the original volume. To remove small molecular impurities (e.g., soluble proteins, peptides, and residual medium components), 3–5 volumes of pre-chilled-phosphate-buffered-saline (PBS, pH 7.4) were added to the sample during concentration for continuous diafiltration. The diafiltered concentrate was collected as the purified EV preparation. The entire procedure was carried out at 4 °C to maximally preserve the integrity and biological activity of the EVs.

### 2.2 Characterization of TH⁺/FOXA2⁺ cell-derived EVs

#### 2.2.1 Transmission Electron Microscopy (TEM)

The purified EV samples were gently resuspended in an appropriate volume of phosphate-buffered saline (PBS, pH 7.4) to obtain a homogeneous suspension without obvious aggregation. Negative staining was used to prepare samples for transmission electron microscopy (TEM) as follows: 10–20 μL of the EV suspension was dropped onto a 200-mesh Formvar-coated copper grid (AZH200, Zhongjing Keyi) and allowed to stand at room temperature for 10 min to enable natural sedimentation and attachment of EVs onto the film; excess liquid was carefully removed with filter paper, and the grid was air-dried at room temperature for approximately 2 min. Then, 20 μL of uranyl acetate staining solution (2%, w/v) was dropped onto the copper grid, and the grid was stained for 5 min at room temperature in the dark (uranyl acetate creates electron density contrast with biological samples, rendering the background dark while leaving the EV vesicle structures bright and clear; when phosphotungstic acid (2%, pH 7.0) was used as an alternative staining solution, the staining time was reduced to 3 min). Excess staining solution was carefully removed from the edge of the copper grid using filter paper, and the grid was placed under an incandescent lamp (approximately 40 W, at a distance of 15–20 cm) to air-dry for an additional 10–15 min until completely dry. The prepared grid was mounted onto the sample holder and placed into a transmission electron microscope (JEOL JEM-1400); observations were performed at an accelerating voltage of 80–100 kV, and images were captured at magnifications of 20,000–50,000×, selecting areas with clearly defined edges and uniform distribution. Typical EVs exhibited cup-shaped, saucer-shaped, or spherical vesicular structures with diameters of approximately 50–150 nm, against a background of high electron density, with clearly outlined vesicle membranes.

#### 2.2.2 Nanoparticle Tracking Analysis (NTA)

Measurements were performed using a Malvern Panalytical NanoSight Pro system in scattered light mode. Prior to detection, the sample chamber was rinsed with ultrapure water to ensure that the number of residual particles in the field of view was fewer than 5. The samples were stored at 4 °C and diluted to an appropriate concentration with DPBS immediately before measurement, mixed thoroughly, and loaded onto the instrument. The injection volume was no less than 500 μL, which was introduced into the sample chamber via a syringe pump. The measurement parameters were set in the software as follows: 750 frames per capture (which could be increased to 1500 frames for low-concentration samples), three measurements per sample, and the optimized distribution (FTLA) selected as the result type. The focus and brightness were manually adjusted to ensure that the number of particles per field of view ranged from 30 to 80 (except for samples where no EVs were expected). The system automatically performed data processing and output the average particle size and concentration. The acceptance criterion was that the difference between the highest and lowest concentrations across the three measurements did not exceed a factor of two, and no overexposed large particles appeared during the acquisition; otherwise, the measurement was repeated. This method effectively evaluates the size distribution and particle concentration of the sample.

#### 2.2.3 Western Blotting (WB)

Purified EV samples were rapidly thawed at 37 °C, mixed with an appropriate volume of RIPA lysis buffer, and lysed on ice for 30 min with intermittent pipetting to ensure thorough mixing. The lysates were then centrifuged at 12,000×g for 5 min at 4 °C, and the supernatants were collected as total EV protein. Protein concentrations were determined using a BCA protein assay kit (Thermo, A55865): EV protein samples were diluted 10-fold with ultrapure water; 25 μL of each standard protein solution at different concentrations, diluted samples, and blank control were added to a 96-well plate (all in duplicate); 200 μL of BCA working solution (Solution A:Solution B = 50:1) was added to each well; the plate was gently mixed and incubated at 37 °C in the dark for 30 min; absorbance was measured at 562 nm using a microplate reader, and a standard curve (R² ≥ 0.98) was plotted to calculate the protein concentration of each sample. EV protein (at least 10 μg per well) was mixed with 4× LDS sample buffer and reducing agent, heated at 70 °C for 10 min for denaturation, and then separated using 4%–12% Bis-Tris precast gels in MES electrophoresis buffer (1×) containing antioxidant agent at a constant voltage of 200 V for 35 min. After electrophoresis, the transfer buffer was prepared by mixing ultrapure water, absolute ethanol, and rapid transfer solution at a ratio of 7:2:1. PVDF membranes were activated and assembled into transfer cassettes in the order of sponge-filter paper-gel-PVDF membrane-filter paper-sponge (with the membrane positioned toward the anode), placed into the transfer tank with an ice pack, and transferred at a constant current of 400 mA for 25 min. After transfer, the PVDF membrane was blocked in blocking buffer on a shaker for 1 h at room temperature, washed three times with 1× TBST for 5 min each. Primary antibodies were diluted in primary antibody dilution buffer as follows: ALIX (1:1000), TSG101 (1:1000), CD9 (1:1000), CD63 (1:1000), CD81 (1:1000), Calnexin (1:20000, negative marker), and GAPDH (1:10000, loading control). The membrane was incubated with primary antibodies overnight at 4 °C on a shaker, washed three times with TBST, and then incubated with HRP-conjugated anti-rabbit secondary antibody (1:2000) for 1 h at room temperature, followed by three TBST washes. Finally, ECL chemiluminescent substrate (equal volume mixing of Solution A and B) was prepared, and the membrane was placed in a chemiluminescence imaging system, covered with the substrate, and exposed for imaging. Based on the marker positions, positive EV markers (ALIX, TSG101, CD9, CD63, CD81) were considered positive if the expected bands appeared, while the absence of a Calnexin band indicated no organelle contamination.

### 2.3 Animal Models and Experimental Groups

The animals used in this study were 6-month-old male A53T transgenic mice (PD-A53T) on a C57BL/6J background, carrying the human α-synuclein gene (SNCA A53T) under the control of the human prion protein promoter (Prnp) to recapitulate the pathological features of PD. Age-matched littermate wild-type C57BL/6J mice served as controls. Qualified mice were sorted in descending order of fall latency and then assigned to different groups using an equidistant sampling method as follows: control group (n = 6, wild-type, DPBS), model group (n = 6, A53T, DPBS), TH⁺/FOXA2⁺ cell-derived EVs high-dose group (n = 7, A53T, 1×10¹¹ particles/mL), TH⁺/FOXA2⁺ cell-derived EVs medium-dose group (n = 6, A53T, 2.5×10¹⁰ particles/mL), and TH⁺/FOXA2⁺ cell-derived EVs low-dose group (n = 6, A53T, 1.25×10¹⁰ particles/mL). All groups received intranasal administration, with 10 μL applied to each nostril (total 20 μL per mouse). The dosing frequency was twice daily for the first three days, followed by once daily thereafter, for a total of 28 consecutive days. During administration, mice were gently restrained, and after intranasal instillation, the position was maintained for 10 seconds to ensure absorption.

The rat model used male Sprague–Dawley-rats weighing 220–250 g, with lesions targeted to the medial forebrain bundle. The injection protocol was as follows: 6-OHDA-at a concentration of 4 μg/μL, injection volume of 2–4 μL, followed by a postoperative observation period of 1–4 weeks to allow lesion stabilization before behavioral testing. After successful modeling, rats were randomly divided into 6 groups (n = 15 rats p-er group): sham-operated group, model group, TH⁺/FOXA2⁺ cell-derived EVs high-, medium-, and low-dose groups (1×10¹¹, 2.5×10¹⁰, and 1.25×10¹⁰ particles/mL, respectively; 80 μL per dose) and a Madopar positive control group (50 mg/kg; 0.4 mL per dose). The dosing regimen was twice daily (BID) for the first 3 days, followed by once daily (QD) for 56 days. All animals were purchased from Jiangsu Huachuang Xinnuo (Production License-No. SCXK (Su) 2020-0009; Quality Certificate No. 2025081201). The experiments were conducted in an SPF animal facility, with facility management following the nationa-l standard GB 14925-2010. The environmental temperature was maintained at 21 °C, relative humidity at 40–70%, under a 12 h/12 h light/dark cycle. Animal use was approved by the Science and Technology Commission of Shanghai Municipality (License No. S-YXK (Shanghai) 2024-0009). All procedures were reviewed and approved by the Institutional Animal Care and Use Committee, complying with animal welfare requirements.

### 2.4 Behavioral Studies

Apomorphine-induced-rotational behavior was used to evaluate the recovery of neurological function in Parkinson’s disease model rats. All rats were tested before administration and on days 7, 14, 28, 42, and 56 after the start of treatment. Following subcutaneous injection of apomorphine at a dose of 0.5 mg/kg, each rat was immediately placed into a custom-built-rotational behavior test chamber (30 cm in diameter, 35 cm in height), and the number of rotations toward the intact side (left side) was recorded over a 30-min-period. The test procedure was video-recorded-and rotational behavior was automatically analyzed. Environmental conditions were kept consistent before testing to ensure data reliability.

The rotarod test was used to evaluate motor coordination and balance. All mice were tested before administration (day 0) and on days 3, 7, 14, 21, and 28 after treatment, while all rats were tested before administration and on days 7, 14, 28, 42, and 56 after treatment. Adaptation training was performed for three days prior to the test, with two sessions per day (4 h apart); the rotation speed was linearly accelerated from 4 rpm to 10 rpm within 2 min, and each training session lasted 5 min. During the formal test, the rotarod had a diameter of 3 cm, and the rotation speed started at 4 rpm and was linearly accelerated to 40 rpm within 2 min, with a maximum test duration of 300 s. Each animal was tested three consecutive times with an interval of at least 30 min between trials, and the latency to fall (the time from the start of rotation until the animal fell) was recorded. The average of the three trials was taken as the final result. The instrument was calibrated before testing, and environmental conditions (lighting, noise, etc.) were kept consistent to ensure data reliability.

The hanging test was used to evaluate muscle strength and motor coordination. The test time points were synchronized with those of the rotarod test. The experimental apparatus consisted of a wire mesh grid (pore size 0.5 cm) fixed horizontally at a height of 40 cm above the floor. During the test, the mouse was gently placed onto the center of the grid, after which the grid was quickly inverted so that the mouse hung upside down by grasping the mesh with its limbs; timing started immediately, and the latency to fall was recorded, with a maximum test duration of 60 s. Each mouse was tested three consecutive times with an interval of 10 min between trials. Based on the latency to fall, scores were assigned as follows: 0–10 s = 1 point, 11–30 s = 2 points, 31–60 s = 3 points, and >60 s = 4 points. The average score of the three trials was calculated as the final result for each time point.

### 2.5 Imaging Studies

Magnetic resonance imaging (MRI) and PET/CT-were used to evaluate brain structure and dopamine synthesis enzyme activity in Parkinson’s disease model rats. All rats underwent MRI scanning on day 57 after the completion of treatment, with anesthesia maintained using 1.5% isoflurane. In addition, dynamic PET/CT-scanning was performed at the end of week 8, and the standardized uptake value ratio (SUVr) of the striatum to the cerebellum was calculated to assess dopamine synthesis enzyme activity. Quantification was based on n = 3 biologically independent animals per group for MRI and n = 2 biologically independent animals per group for PET/CT. The instruments were calibrated before testing, and environmental conditions (lighting, noise, etc.) were kept consistent to ensure data reliability.

### 2.6 Generation of Midbrain Organoids (MBOs) and Establishment of the PD Model

To generate midbrain organoids, the STEMdiff™ Midbrain Organoid Differentiation Kit (Catalog #100-1096, STEMCELL Technologies) was used according to the manufacturer’s instructions. Briefly, human pluripotent stem cells (hPSCs) were dissociated into single cells and seeded into AggreWell™ 800 microwell plates (STEMCELL Technologies) at a density of 7,000 cells per microwell to form embryoid bodies. After 5 days, the embryoid bodies were transferred to suspension culture and patterned in STEMdiff™ Midbrain Organoid Differentiation Medium. On day 6, the organoids were transferred to STEMdiff™ Neural Organoid Maintenance Medium for long-term-culture (>50 days) to allow further maturation. The generated organoids were characterized for the midbrain-specific-marker FOXA2 and the dopaminergic neuron marker TH. To confirm MBO identity, immunofluorescence staining was performed for the floor-plate/midbrain progenitor markers FOXA2, LMX1A and CORIN, the dopaminergic neuron markers TH, NURR1 (NR4A2) and DAT, the pan-neuronal marker TUJ1 and the neural progenitor marker SOX2, together with the pluripotency marker OCT4 as a negative control.

For the PD model, mature midbrain organoids cultured for 60 days were transferred to 24-well-low-attachment-plates (3–5 organoids per well) and incubated with the above midbrain differentiation medium containing 6-hydroxydopamine-(6-OHDA-) at a final concentration of 300μM for 48 h in a 37 °C, 5% CO₂ cell culture incubator. After treatment, the 6-OHDA-containing--medium was aspirated, and the organoids were gently washed twice with pre-chilled-PBS for 5 min each time, then returned to normal midbrain differentiation medium for an additional 24–72 h to establish a stable *in vitro* model of PD. For EV intervention, two dosing regimens were compared. In the TH⁺/FOXA2⁺ cell-derived EVs-1 group (post-modeling treatment), EVs were not present during 6-OHDA exposure: TH⁺/FOXA2⁺ cell-derived EVs were added at 5×10¹⁰ particles/mL only after the 48-h 6-OHDA modeling period had been completed and the 6-OHDA-containing medium had been removed. In the TH⁺/FOXA2⁺ cell-derived EVs-2 group (concurrent treatment), the same concentration of TH⁺/FOXA2⁺ cell-derived EVs (5×10¹⁰ particles/mL) was added together with 300 μM 6-OHDA at the onset of modeling, so that modeling and treatment proceeded simultaneously. After the initial 48 h, both groups were maintained in normal midbrain differentiation medium containing 5×10¹⁰ particles/mL of TH⁺/FOXA2⁺ cell-derived EVs for an additional 7 days.

### 2.7 Histological Analysis

In this study, brain tissue sections from A53T transgenic mice and MBO models were used for PD--related pathological analysis. Tissues were harvested 28 days after treatment. Mouse brain tissue was subjected to cardiac perfusion followed by fixation with 4% paraformaldehyde. Whole brains were then dehydrated through a graded ethanol series, embedded in paraffin, and serially sectioned at a thickness of 4 μm, which were mounted onto adhesion slides for subsequent use. Human midbrain organoids were fixed with 4% paraformaldehyde after culture, dehydrated in sucrose, embedded in OCT compound, and sectioned at a thickness of 10 μm using a cryostat. To evaluate histomorphological changes, brain sections were stained with hematoxylin and eosin (H&E) for observation of tissue architecture, and Nissl staining was performed to assess neuronal survival. Immunohistochemical staining was used to detect the distribution of TH-positive neurons and fiber density, while α-sy-n staining was employed to observe pathological protein aggregation. For midbrain organoids, immunofluorescence double-labelin-g was performed using MAP2/TH co-stainin-g to evaluate the maturation and dendritic integrity of dopaminergic neurons, and GFAP/TUJ1 co-stainin-g to observe the relationship between astrocyte activation and neuronal architecture. All stained images were captured using a microscope and quantitatively analyzed with ImageJ software. Quantitative analyses were performed on n = 3 biologically independent animals per group.

### 2.8 Flow Cytometry Analysis

In this study, flow cytometry was used to detect the surface marker SSEA4 and the intracellular marker Nanog in iPSCs, as well as the intracellular markers TH, FoxA2, and β-III Tubulin in dopaminergic neurons. First, a single-cell-suspension was prepared: neural cell clusters were digested with a solution containing papain and DNase I at 37 °C for 8–12 min, gently pipetted to disperse, then mixed with stop solution and filtered through a 100μm cell strainer, and viable cells were counted. A total of 1×10⁷ cells were collected, centrifuged, and resuspended. For viability staining, Zombie Aqua™ dye was added and incubated for 15–30 min at 4 °C in the dark, followed by washing and resuspension. Human TruStain FcX was added to block non-specific binding for 5–10 min at room temperature. After aliquotting, cells for surface marker SSEA4 were stained with PE-conjugated-antibody or the corresponding isotype control, incubated for 15–20 min at 2–8 °C in the dark, washed, and resuspended for analysis. For intracellular staining, cells were first fixed and permeabilized: 1× Fix/Perm Buffer was added and incubated for 40–50 min at 2–8 °C in the dark, followed by washing with Perm/Wash Buffer. Subsequently, cells were stained with Alexa Fluor® 647 anti-Nanog-, TH-PE-, FoxA2 Alexa Fluor® 488, and APC anti-Tubulin-β-3 antibodies, along with their respective isotype controls, incubated for 40–50 min at 2–8 °C in the dark, washed, and resuspended. Analysis was performed on a BD FACS Canto II flow cytometer. The main cell population P1 was first gated, followed by single cells and viable cells. The negative gate was set using isotype controls (positive rate controlled below 0.4%), and the positive percentages of each marker were determined. This method enables standardized multi-marker-detection of cells.

### 2.9 Toxicological Evaluation

In this study, 30 SD rats (15 per sex) were randomly divided into 3 groups (5 rats/group/sex). Group 1 served as the vehicle control group and received PBS (pH 7.4). Groups 2 and 3 were the TH⁺/FOXA2⁺ cell-derived EVs-LD-and TH⁺/FOXA2⁺ cell-derived EVs-HD-groups, which received the test article (TH⁺/FOXA2⁺ cell-derived EVs) at doses of 1×10¹⁰ and 4×10¹⁰ particles per animal, respectively. All animals were administered intranasally once daily for 30 consecutive days. In addition, six cynomolgus monkeys were randomly divided into 3 groups (one male and one female per group). Group 1 received PBS (pH 7.4) as the vehicle control, while Groups 2 and 3 received TH⁺/FOXA2⁺ cell-derived EVs at doses of 2×10¹⁰ and 8×10¹⁰ particles per animal as the low-dose-and high-dose-groups, respectively. The animals received repeated intranasal spray administration (using a single-use-nasal atomization device; Registration No. SuXieZhun 20192080271---) once daily for 30 consecutive days.

During the study period, clinical observations (including general clinical observation, detailed clinical observation, local observation at the administration site (both nostrils of each animal), body weight, food consumption, body temperature, clinical pathology (hematology, coagulation function, serum biochemistry), and gross necropsy) were performed. All animals were euthanized on day 31, and scheduled animals underwent systematic necropsy and gross observation. Routine histopathological examination was performed on protocol-specifie-d tissues, primarily those related to drug absorption and important organs, including brain, nasal cavity, olfactory bulb, heart, liver, lung, and spleen.

### 2.10 Transcriptomic Sequencing (the 6-OHDA-Induced MBO PD Model)

Library Construction and Sequencing: Total RNA was extracted, and mRNAs with poly(A) tails-were enriched using Oligo(dT) magnetic beads. The enriched RNA was fragmented with fragmentation buffer, and reverse transcription was performed using random N6 primers to synthesize the second strand of cDNA, generating double--stranded DNA. The doubl-e-stranded DNA was subjected to end repair, 5′ phosphorylation, and addition of a single ‘A’ overhang at the 3′ end, followed by ligation with a bubble-shap-ed adapter having a protruding ‘T’ at the 3′ end. The ligation products were amplified by PCR using specific primers. The PCR products were heat-denatur-ed into single strands and circularized via a bridge primer to obtain a single-stra-nd circular DNA library. After library quality control, paired-e-nd 150 bp (PE150) sequencing was performed on the DNBSEQ platform.

Data Analysis: The raw sequencing data (Raw data) were quality-controlled- and filtered using SOAPnuke (v1.5.6). Reads containing adapter contamination, unknown bases (N) exceeding 1%, or low-quality-bases (Q < 15) accounting for more than 20% of the read were removed to obtain Clean data. The Clean data were aligned to the reference gene set using Bowtie2 (v2.3.4.3), and gene expression quantification was performed using RSEM (v1.3.1). A sample-to-sample--gene expression clustering heatmap was generated using pheatmap (v1.0.12). Differential expression analysis between groups was performed using DESeq2 (v1.4.5) with a screening threshold of Q value < 0.05 and fold change > 1.5. GO, KEGG, and Reactome enrichment analyses of differentially expressed genes were carried out using clusterProfiler (v4.14.0), with a p-value-< 0.05 set as the threshold for significant enrichment.

### 2.11 Proteomics (TH⁺/FOXA2⁺ cell-derived EVs and the 6-OHDA-Induced MBO PD Model)

Protein Digestion: A 100μg protein sample was loaded into a 10 kDa ultrafiltration tube. When processing multiple samples simultaneously, the volumes of all samples were equalized to allow centrifugal balance. The volume was adjusted with a buffer that was identical to the sample dissolution buffer. If all sample volumes were below 100 μL, the final volume was made up to 100μL; if any sample volume exceeded 100μL, the volume of all samples was adjusted to match that of the largest one. The tube was centrifuged at 20 °C and 12,000×g for 20 min until all protein solution had been spun to the bottom of the collection tube. Then, 100μL of 0.5 M TEAB was added, and the tube was centrifuged again at 20 °C and 12,000×g for 20 min; this step was repeated three times. A new collection tube was used. Trypsin was added at an enzyme-to-protein--ratio of 1:20, and the mixture was incubated at 37 °C for 4 h. After the reaction, the tube was centrifuged at 20 °C and 12,000×g for 15 min, and the digested peptide solution was collected at the bottom of the tube. An additional 100μL of 0.5 M TEAB was added to the ultrafiltration tube, followed by centrifugation at 20 °C and 12,000×g for 15 min. The collected peptide digests were pooled and lyophilized to dryness.

Mass Spectrometry Detection: The dried peptide samples were reconstituted in mobile phase A (H₂O containing 0.1% FA) and centrifuged at 20,000×g for 10 min, after which the supernatant was injected for analysis. Separation was performed on a Thermo Vanquish Neo liquid chromatography system. The sample was loaded onto an EASY-Spray-™ HPLC column (150μm × 15 cm, Thermo, USA) and separated at a flow rate of 2.5μL/min using the following effective gradient: 0–4 min, mobile phase B (80% ACN, 0.1% FA) linearly increased from 4% to 25%; 4–5.8 min, mobile phase B increased linearly from 25% to 35%; 5.8–6.2 min, mobile phase B increased from 35% to 99% with the flow rate increased to 3μL/min; 6.2–6.9 min, 99% mobile phase B. The outlet of the nano-LC separation was directly connected to the mass spectrometer. The LC-separated-peptides were ionized using a nanoESI source and then introduced into a tandem mass spectrometer (Astral, Thermo, USA) operated in data-independent-acquisition (DIA) mode. The main parameters were set as follows: ion source voltage, 1.8 kV; MS1 scan range, 380–980 m/z; resolution, 240,000; maximum injection time (MIT), 5ms. The mass range of 380–980 m/z was divided into 300 windows for sequential windowed fragmentation and signal acquisition. Fragmentation was performed using HCD with a normalized collision energy (NCE) of 25, MIT of 3 ms, and fragment ions were detected in the Astral analyzer. The AGC target was set to 500% for MS1 and 500% for MS2.

Data Analysis: The raw mass spectrometry data were analyzed using DIA-NN (v2.2.0) software. Peptide and protein identification and quantification were performed by searching the spectral library against the human proteome database (UniProt Proteome ID: UP000005640). Differential expression analysis was carried out using limma, with significantly differentially expressed proteins screened at a threshold of fold change > 1.5 and p-value-< 0.05.

### 2.12 Single-Cell RNA Sequencing (scRNA-seq) of the 6-OHDA-Induced MBO PD Model

Library Preparation and Sequencing: The cell suspension was stained with 0.4% trypan blue, examined under a microscope, and the viability was calculated. Samples with viability greater than 80% proceeded to library preparation. Library preparation was performed using the DNBelab C Series High-throughput-Single-cell-RNA Library Preparation Kit Set V3.0 (MGI, China). The prepared single-cell-suspension, oil, and beads were sequentially loaded into the C4 scRNA cartridge and apparatus to generate droplets, within which cell lysis and mRNA capture by beads occurred. Subsequently, the droplets were broken by a vacuum pump to release the mRNA-bead-complexes. cDNA synthesis was carried out at appropriate temperatures, followed by amplification and purification of cDNA and oligo products. The concentration and fragment size distribution of the cDNA and oligo products were checked to ensure they were within acceptable ranges. The oligo library was constructed, then amplified, indexed, and purified for circularization. The cDNA products were subjected to fragmentation, end repair, and A-tailing-. The cDNA was ligated to adapters at an appropriate temperature for a set period, then purified. Adapter-ligated-cDNA was amplified by PCR and purified. The cDNA and oligo products were separately denatured into single strands. After denaturation, a circularization reaction system was prepared and the reaction program was run to obtain single-strand-circular products, followed by digestion of non-circularized-linear DNA molecules. Single--stranded circular DNA molecules were replicated via rolling circle amplification to generate DNA nanoballs (DNBs) containing multiple copies of the DNA. High-intens-ity DNA nanoball chip technology was then used to load sufficient DNA into the mesh-l-ike wells on the chip, and sequencing was performed using combinatorial Probe-Anc-hor Synthesis (cPAS). The cDNA library was sequenced in PE47+100 mode, and the oligo library in PE32+42 mode.

Data Analysis: Raw sequencing data from each sample were processed using DNBelab_C4scRNA (v1.0.1) to generate the raw gene expression matrix, and downstream analysis was performed using the R package Seurat (v5.3.0). Quality control was applied to retain only cells with UMI counts between 500 and 25,000 (500 < nCount < 25,000), number of detected genes between 200 and 10,000 (200 < nFeature < 10,000), and mitochondrial gene percentage below 10% (percent.mt < 10%). Meanwhile, DoubletFinder (v2.0.6) was used to identify and remove potential doublets. After quality control, a total of 50,729 high-quality-cells were obtained for subsequent analysis. Data were normalized using the SCTransform method, the top 3,000 highly variable genes were selected, and the percentage of mitochondrial gene expression was regressed out as a covariate to eliminate technical variation. Principal component analysis (PCA) was then performed, and the first 30 principal components (PCs) were selected for UMAP visualization and subsequent clustering. To eliminate batch effects across different samples, an integration method based on canonical correlation analysis (CCA) was applied to correct the data. Based on the integrated data, unsupervised clustering analysis was performed using the FindNeighbors and FindClusters functions with the resolution parameter set to 1.0, resulting in the identification of 24 cell clusters. To identify specific marker genes for each cluster, the FindAllMarkers function in the Seurat package was used with Wilcoxon rank-sum-test. Differentially expressed genes (DEGs) were screened using the following criteria: the gene was upregulated only in the target cluster (only.pos = TRUE), log2 fold change > 0.25 (logfc.threshold > 0.25), expressed in at least 25% of cells in the target cluster (min.pct > 0.25), and p-value-< 0.05. Cell type annotation was performed for each cluster by combining canonical marker genes and the identified DEGs. Finally, cell-cell-communication analysis and visualization among cell subpopulations were performed using the CellChat (v1.6.1) package.

### 2.13 Small RNA Sequencing (TH⁺/FOXA2⁺ cell-derived EVs)

Library Preparation and Sequencing: Library construction was performed using the MGIEasy Small RNA Library Preparation Kit (BGI-Shenzhen, China). An appropriate amount of RNA sample was subjected to sequential 3′ and 5′ adapter ligation, reverse transcription, and PCR amplification. Subsequently, the target bands were recovered by polyacrylamide gel electrophoresis (PAGE). The PCR products were denatured into single-strand-ed DNA, circularized, and the non-circularized-linear DNA molecules were digested to obtain the final library. After denaturing the library into single strands, DNA nanoballs (DNBs) were generated by amplification using phi29 polymerase and loaded onto the sequencing chip. Sequencing was performed on the G400 sequencer (BGI-Shenzhen, China) using combinatorial Probe-Anchor-Synthesis (cPAS) technology in SE50 mode.

Data Analysis: Raw sequencing data were filtered to remove low-quality-sequences (containing ≥4 bases with quality value <10 or ≥6 bases with quality value <13), sequences with 5′ adapter contamination, missing 3′ adapters, sequences without inserts, sequences with poly(A) tails-, and sequences shorter than 15nt, yielding high-quality-data (Clean data). The Clean data were aligned to the reference genome and the miRBase database using Bowtie2. Based on these alignments, miRDeep2 (v0.1.3) was used to predict novel miRNAs, and known miRNAs were identified based on the alignment results against miRBase. In addition, cmsearch (v1.1.2) was used to align against the Rfam database to achieve comprehensive annotation of miRNAs and other non-coding-RNAs. Unique Molecular Identifiers (UMIs) are short sequences of 8–10nt that are ligated to cDNA molecules at an early stage of library construction to label each molecule in the original sample. Counting the number of distinct UMI species effectively eliminates quantification bias introduced by PCR amplification preference, thereby ensuring accurate small RNA quantification. Expression levels were calculated using the formula: Expression = (C × 1,000,000) / T, where C is the number of UMIs aligned to a given miRNA, and T is the total number of UMIs from clean reads of the sample. Differential expression analysis was performed using DESeq2. The thresholds for significantly differentially expressed miRNAs were set as fold change > 1.5 and q-value-< 0.05. Interaction pairs between differentially expressed miRNAs and target genes were screened based on the miRTarBase database (v10.0), and the miRNA–mRNA-regulatory network was constructed and visualized using Cytoscape software.

### 2.14 Statistical Analysis

Statistical analyses were performed using GraphPad Prism 8.4 software. Data were presented as mean ± standard deviation (mean ± SD). The Shapiro–Wilk-test was used to assess the normality of data distribution, and Levene’s test was used to evaluate homogeneity of variances among groups. For comparisons between two groups, Student’s t-test-was used for normally distributed data, otherwise the Mann–Whitney-U test was applied. For comparisons among multiple groups, one-way-analysis of variance (ANOVA) followed by Tukey’s multiple comparison test was performed. For repeated measures data (e.g., behavioral outcomes), repeated measures ANOVA or a mixed-effects-model was used to analyze the interaction between group and time, and comparisons among groups at each time point were performed using one-way-ANOVA followed by Tukey’s multiple comparison test. A p-value-≤ 0.05 was considered statistically significant. Unless otherwise stated, “n” denotes the number of biologically independent samples (biological replicates), i.e., independent animals or independent organoid/EV preparations; technical replicates were averaged within each biological replicate and were not treated as independent observations. Every figure legend indicates the corresponding n and whether biological replicates were used.

## 3. Results

### 3.1 Differentiation and Characterization of TH⁺/FOXA2⁺ Neural Cells and EVs

We first established a workflow for differentiating human iPSCs into TH⁺/FOXA2⁺ neural cells (Fig. 1A, 1B). Flow cytometric analysis on differentiation days 24, 30, and 60 showed that the positive rate of the undifferentiated marker SSEA4 was extremely low (<2%), while the positive rates of the key dopaminergic neuron transcription factors FOXA2 and TH both exceeded 90% (Fig. 1C), indicating highly efficient specification toward a homogeneous TH⁺/FOXA2⁺ neural cell population. Given the persistent co-expression of the floor-plate factor FOXA2, this population is best regarded as midbrain floor-plate-derived TH⁺/FOXA2⁺ neural cells at an intermediate differentiation stage, rather than as fully mature, terminally differentiated midbrain dopaminergic neurons. EVs were extracted from the supernatants collected every two days between days 30 and 60 of differentiation and were designated as TH⁺/FOXA2⁺ cell-derived EVs. Transmission electron microscopy (TEM) revealed that TH⁺/FOXA2⁺ cell-derived EVs exhibited typical disc-shaped-bilayer membrane structures (Fig. 1D); nanoparticle tracking analysis (NTA) showed a major size peak at approximately 100 nm (Fig. 1E); western blotting detected the positive EV markers CD9, CD81, TSG101, and ALIX, while the endoplasmic reticulum protein Calnexin was negative (Fig. 1F), demonstrating that TH⁺/FOXA2⁺ cell-derived EVs were highly pure and free from cellular debris contamination.

**Figure 1.**
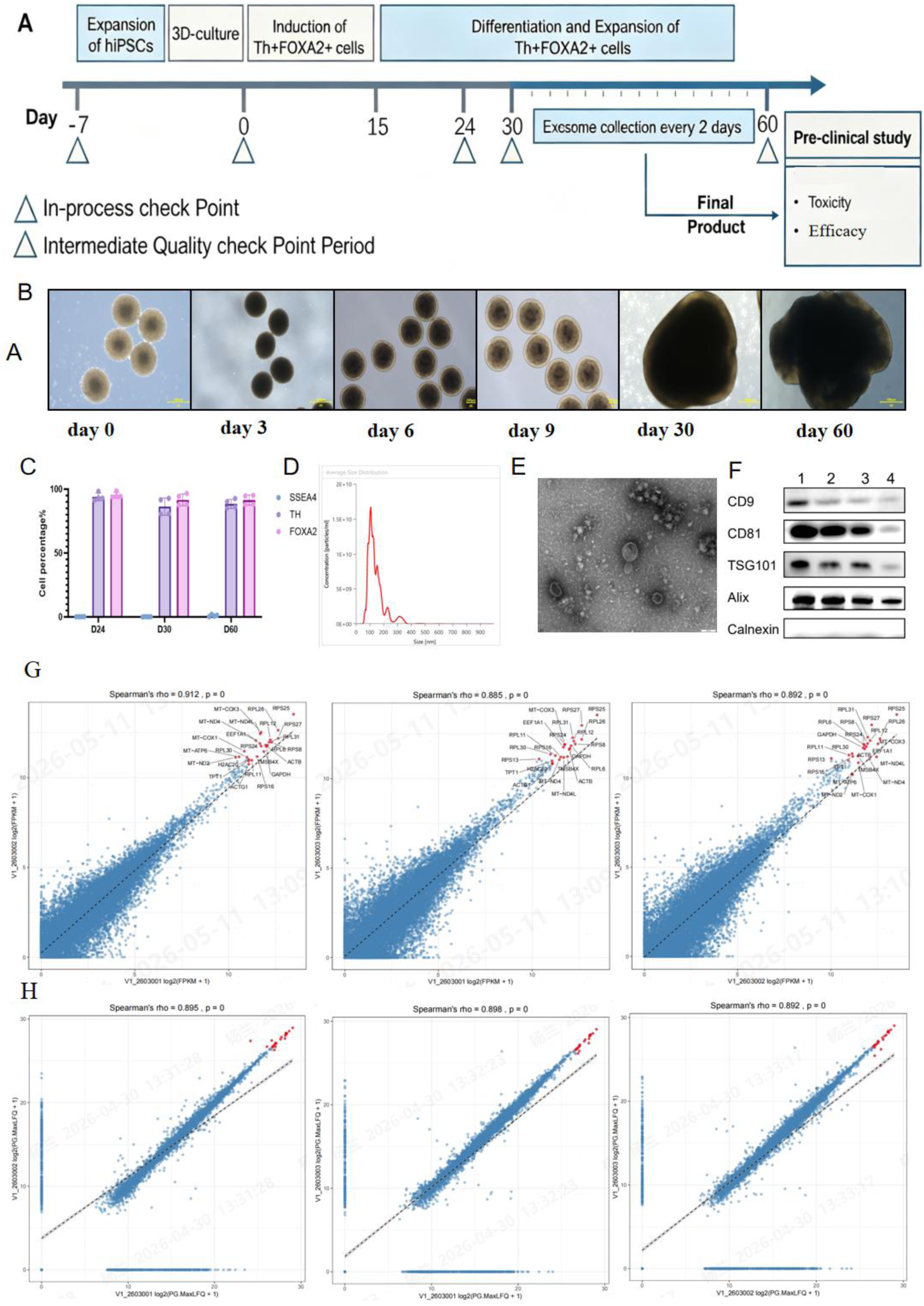
Differentiation of neural cells and characterization of TH⁺/FOXA2⁺ cell-derived EVs. Differentiation of neural cells and characterization of TH⁺/FOXA2⁺ cell-derived EVs.(A) Workflow for the production of TH⁺/FOXA2⁺ cell-derived EVs. (B) Differentiation protocol for neural cells. (C) Flow cytometric analysis of the differentiation efficiency of neural cells (n = 4). (D) Representative transmission electron microscopy (TEM) image showing the disc-shaped-bilayer membrane structure of TH⁺/FOXA2⁺ cell-derived EVs. (E) Nanoparticle tracking analysis (NTA) of the size and concentration of TH⁺/FOXA2⁺ cell-derived EVs. (F) Western blotting (WB) detection of marker proteins of TH⁺/FOXA2⁺ cell-derived EVs. (G)Spearman correlation coefficient of mRNAs of TH⁺/FOXA2⁺ cell-derived EVs across three batches. (H)Spearman correlation coefficient of proteins of TH⁺/FOXA2⁺ cell-derived EVs across three batches. Biological replicates: (C) n = 4 independent differentiation experiments; (D)–(H) n = 3 independent TH⁺/FOXA2⁺ EV batches. (D)–(F) are representative images of n = 3 biologically independent EV preparations.

Transcriptomic and proteomic analyses of TH⁺/FOXA2⁺ cell-derived EVs revealed 5,021 co-expressed-genes at both mRNA and protein levels, which were enriched in multiple factors associated with neural repair and regeneration. Enrichment analysis of the top 100 factors showed GO enrichment in neurodevelopmental processes (midbrain, substantia nigra, and neural nucleus development), protein homeostasis (degradation and autophagy), and energy metabolism; KEGG enrichment was observed in Parkinson’s disease and other neurodegenerative disease pathways, proteasome and phagosome (protein aggregation and degradation pathways), as well as glycolysis and other energy metabolism pathways. To verify the consistency among different batches of EVs, we systematically evaluated their homogeneity. Across three independent batches collected between days 30 and 60, a total of 11,646 co-expressed genes and 4,760 co-expressed proteins were identified, with good inter-batch c-onsistency (Fig. 1G, 1H). The top 20 genes and top 20 proteins were -stably and highly expressed across batches. These results demonstrate tha-t the EVs used in this study exhibit excellent homogeneity at both the protein and mRNA levels.

### 3.2 In Vivo Experiments

#### 3.2.1 TH⁺/FOXA2⁺ cell-derived EVs Ameliorate Motor Dysfunction and Alleviate Neuronal Damage in A53T PD Mice

To evaluate the efficacy of TH⁺/FOXA2⁺ cell-derived EVs in a genetic model of Parkinson’s disease, A53T mice received intranasal administration for 4 weeks (Fig. 2A). The rotarod test primarily assesses the comprehensive ability of the central nervous system to control motor coordination, balance, and motor endurance. In the rotarod test, all TH⁺/FOXA2⁺ cell-derived EVs dose groups exhibited significantly prolonged fall latencies relative to controls from day 7 onward, with the effect persisting through day 28 (Fig. 2B), suggesting that within the tested dose range, TH⁺/FOXA2⁺ cell-derived EVs exerted sustained and stable improvement in motor coordination. The hanging test mainly reflects limb muscle strength and endurance. In the hanging test, all TH⁺/FOXA2⁺ cell-derived EVs dose groups also showed improvement from day 7 (Fig. 2C), indicating that TH⁺/FOXA2⁺ cell-derived EVs improved hanging performance, thereby suggesting a positive effect on muscle strength/endurance.

**Figure 2.**
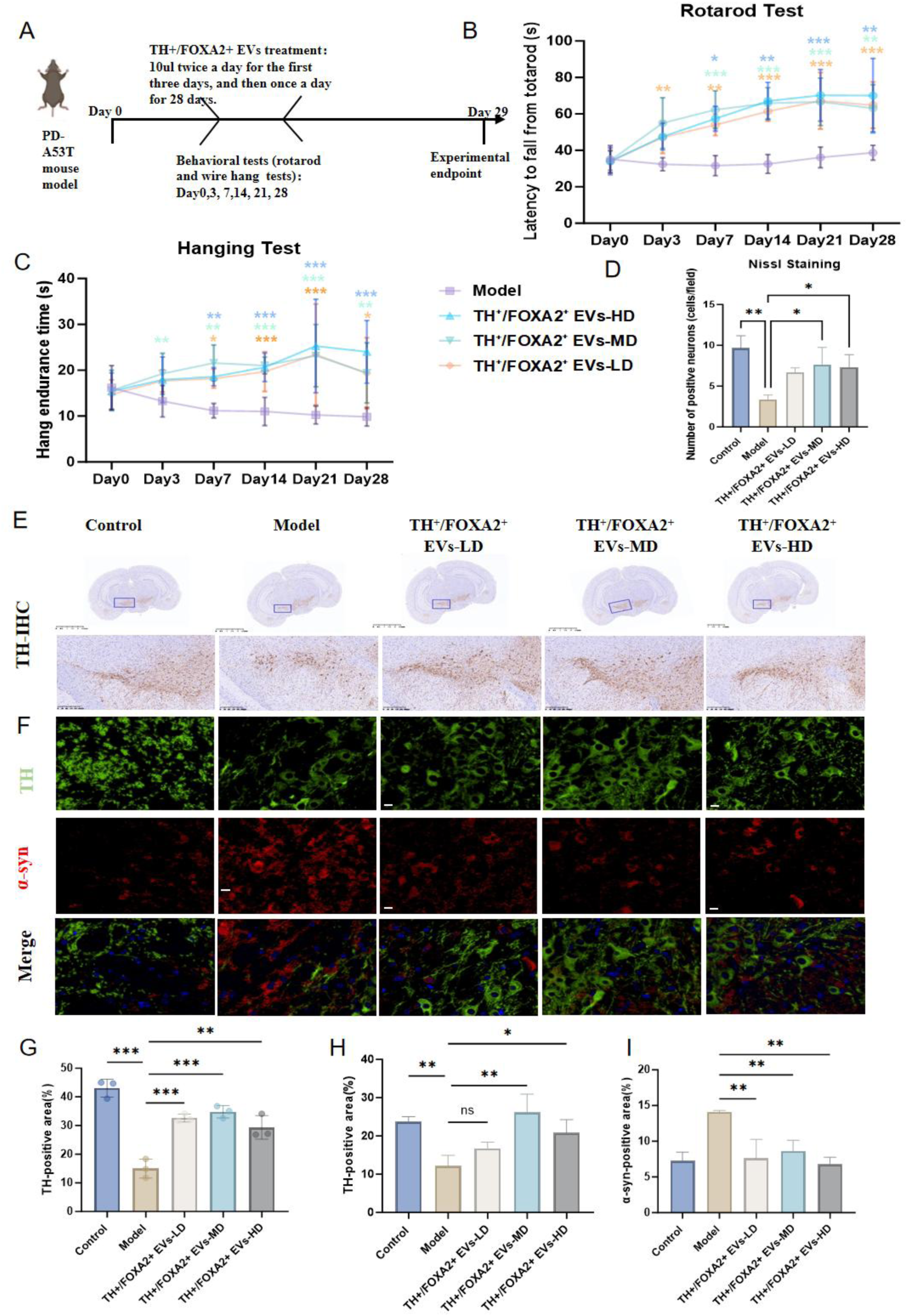
TH⁺/FOXA2⁺ cell-derived EVs Ameliorate Motor Dysfunction and Attenuate Neuronal Damage in the Substantia Nigra of A53T PD Mice. TH⁺/FOXA2⁺ cell-derived EVs improve motor function and alleviate neuronal damage in PD mice.(A) Experimental flowchart of TH⁺/FOXA2⁺ cell-derived EVs treatment in A53T PD-mice. (B) Rotarod test results of PD mice treated with different doses of TH⁺/FOXA2⁺ cell-derived EVs for 4 weeks. (C) Hanging test results of PD mice treated with different doses of TH⁺/FOXA2⁺ cell-derived EVs for 4 weeks. (D) Quantitative analysis of Nissl staining in brain tissues from PD mice treated with different doses of TH⁺/FOXA2⁺ cell-derived EVs for 4 weeks. (E) Immunohistochemical staining of TH in brain tissues from PD mice treated with different doses of TH⁺/FOXA2⁺ cell-derived EVs for 4 weeks. (F) Immunofluorescence staining of TH and α-syn-in brain tissues from PD mice treated with different doses of TH⁺/FOXA2⁺ cell-derived EVs for 4 weeks. (G) Quantitative analysis of TH immunohistochemistry in brain tissues from PD mice treated with different doses of TH⁺/FOXA2⁺ cell-derived EVs for 4 weeks. (H) Quantitative analysis of TH immunofluorescence in brain tissues from PD mice treated with different doses of TH⁺/FOXA2⁺ cell-derived EVs for 4 weeks. (I) Quantitative analysis of α-syn-immunofluorescence in brain tissues from PD mice treated with different doses of TH⁺/FOXA2⁺ cell-derived EVs for 4 weeks. n = 3. Scale bar = 50μm. Data are shown as mean ± SD. ns, not significant; *p<0.05, **p<0.01, ***p<0.001 vs. model group. All quantifications were performed on biologically independent animals (biological replicates), n = 3 per group; (E) and (F) are representative images from n = 3 biologically independent animals per group.

To further evaluate the protective effect of TH⁺/FOXA2⁺ cell-derived EVs on dopaminergic neuron injury in A53T PD mice, systematic histopathological and immunohistochemical analyses were performed on mouse brain tissues. Nissl staining showed that the number of Nissl bodies in the substantia nigra was markedly reduced and the staining was lighter in the model group, indicating impaired neuronal synthetic function; after intervention with TH⁺/FOXA2⁺ cell-derived EVs-HD and EVs-MD groups, the number of Nissl bodies was significantly increased with deeper staining, suggesting restoration of neuronal metabolic activity (Fig. 2D).

Further immunohistochemical and immunofluorescence results showed that the number of TH-positive-neurons and the density of TH-positive-fibers were significantly reduced in the model group, indicating severe loss of dopaminergic neurons and their projecting fibers. In contrast, after treatment with TH⁺/FOXA2⁺ cell-derived EVs-HD- and EVs-MD, the number of surviving TH-posi-tive neurons was significantly increased, and fiber density was markedly restored (Fig. 2E, 2F, 2G, 2H). α-Synuclein (α-syn) staining further revealed abundant α-syn-positive aggregat-es in t-he model group, whereas treatment with different do--ses of TH⁺/FOXA2⁺ cell-derived EVs significantly reduced pathological α-syn protein aggregation (Fig. 2F, 2I). Collectively, these results dem-onstrate that TH⁺/FOXA2⁺ cell-derived EVs effectively alleviate dopaminergic neuron degeneration in the midbrain substantia nigra of A53T PD mice, inhibit α-syn pathological aggregation, and improve neuronal functional status.-

#### 3.2.2 TH⁺/FOXA2⁺ cell-derived EVs Ameliorate Motor Dysfunction and Alleviate Neuronal Damage in the Substantia Nigra of 6-OHDA-Induced PD Rats

Next, we validated the therapeutic efficacy of TH⁺/FOXA2⁺ cell-derived EVs in the 6-OHDA-unilateral lesion rat model (Fig. 3A). In the apomorphine-induced-rotation test, rats in the model group exhibited significant contralateral (intact side) rotations following apomorphine injection, with a markedly increased number of rotations compared to the control group. The TH⁺/FOXA2⁺ cell-derived EVs-HD-group showed a significant reduction in rotations from day 7 onward, achieving an effect comparable to that of Madopar, and by day 28, all TH⁺/FOXA2⁺ cell-derived EVs dose groups exhibited significant therapeutic effects (Fig. 3B). The hanging test and rotarod test further assessed limb muscle strength and motor coordination in rats. Rats in the model group showed a significantly shortened hanging time and a markedly reduced rotarod latency, indicating severe motor impairment. In the hanging test (Fig. 3C), all TH⁺/FOXA2⁺ cell-derived EVs dose groups already showed significant efficacy by day 14, and after day 42, the improvement effects in the TH⁺/FOXA2⁺ cell-derived EVs-MD- and EVs-HD groups were even more pronounced. Similarly, the rotarod test results (Fig. 3D) showed that different doses of TH⁺/FOXA2⁺ cell-derived EVs exerted varying degrees of therapeutic effects as early as day 7; by day 56, the efficacy in the TH⁺/FOXA2⁺ EV-s-MD and EVs-HD groups was significantly better than that in the -low-dose group.

**Figure 3.**
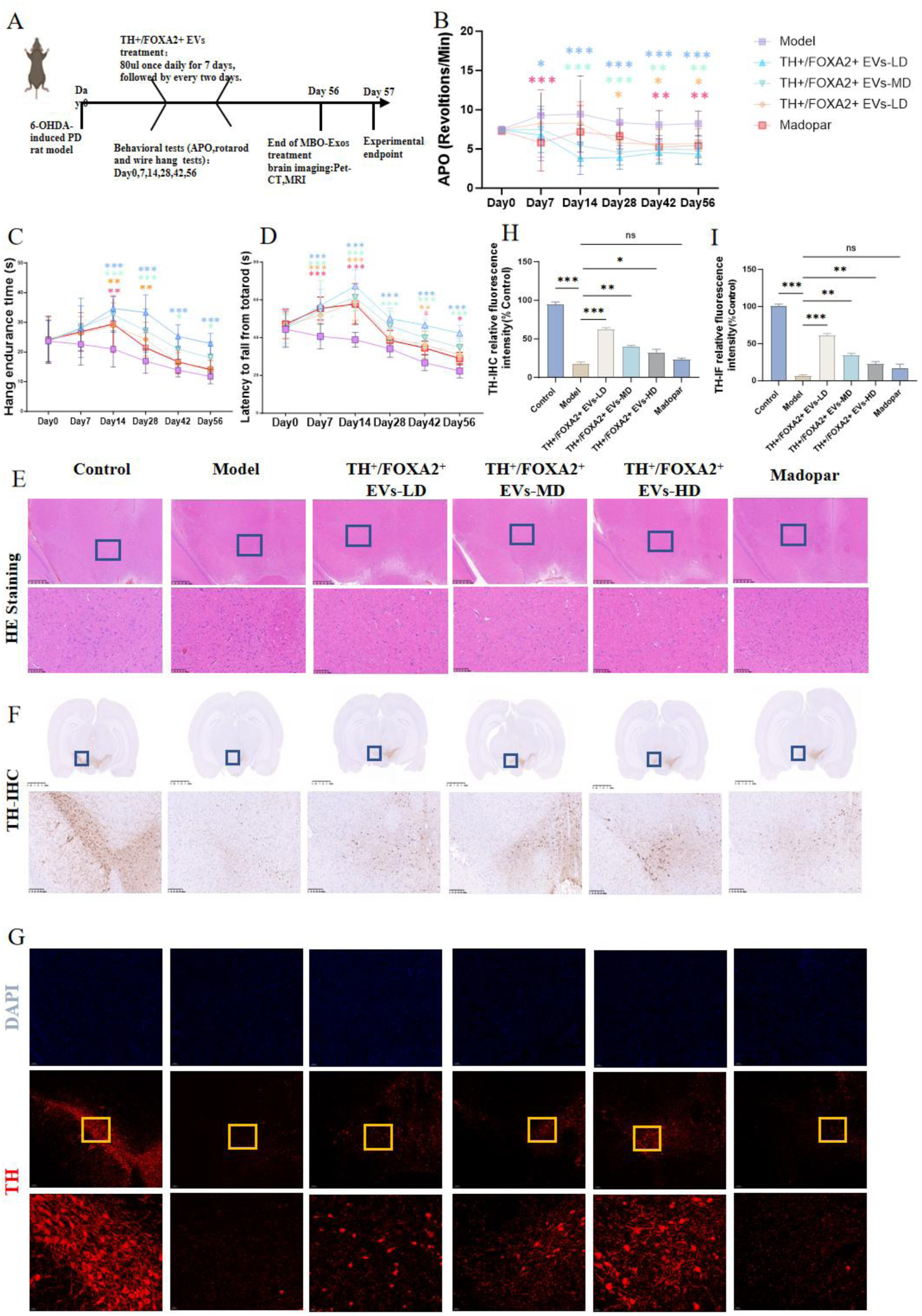
TH⁺/FOXA2⁺ cell-derived EVs improve the locomotor ability and reduce neuronal damage in the substantia nigra of 6-OHDA-induced PD rats. TH⁺/FOXA2⁺ cell-derived EVs improve the locomotor ability and reduce neuronal damage in the substantia nigra of 6-OHDA-induced PD rats.(A) Experimental flowchart of TH⁺/FOXA2⁺ cell-derived EVs treatment in 6-OHDA-PD rats. (B) Apomorphine-induced-rotation test results of PD rats treated with different doses of TH⁺/FOXA2⁺ cell-derived EVs for 8 weeks (n = 15). (C) Hanging test results of PD rats treated with different doses of TH⁺/FOXA2⁺ cell-derived EVs for 8 weeks (n = 15). (D) Rotarod test results of PD rats treated with different doses of TH⁺/FOXA2⁺ cell-derived EVs for 8 weeks (n = 15). (E) H&E staining of brain tissues from PD rats treated with different doses of TH⁺/FOXA2⁺ cell-derived EVs for 8 weeks. (F) Immunohistochemical staining of TH in brain tissues from PD rats treated with different doses of TH⁺/FOXA2⁺ cell-derived EVs for 8 weeks. (G) Immunofluorescence staining of TH in brain tissues from PD rats treated with different doses of TH⁺/FOXA2⁺ cell-derived EVs for 8 weeks. (H) Quantitative analysis of TH immunohistochemistry in brain tissues from PD rats treated with different doses of TH⁺/FOXA2⁺ cell-derived EVs for 8 weeks (n = 3). (I) Quantitative analysis of TH immunofluorescence in brain tissues from PD rats treated with different doses of TH⁺/FOXA2⁺ cell-derived EVs for 8 weeks (n = 3). Scale bar = 50 μm. Data are shown as mean ± SD. ns, not significant; *p<0.05, **p<0.01, ***p<0.001 vs. model group. (E)–(G) are representative images from n = 3 biologically independent animals per group. All n values refer to biologically independent animals (biological replicates); technical replicates were averaged within each animal and were not treated as independent observations.

To further evaluate the protective effect of TH⁺/FOXA2⁺ cell-derived EVs against 6-OHDA-induced--dopaminergic neuron injury in PD rats, systematic histopathological examination was performed on rat brain tissues. H&E staining showed that in the model group, neurons in the lesioned substantia nigra exhibited obvious degenerative changes, including neuronal shrinkage, pyknosis, and hyperchromatic cytoplasm, with disorganized cellular arrangement. After treatment with TH⁺/FOXA2⁺ cell-derived EVs, the morphology of neurons in the substantia nigra improved and degenerative changes were alleviated (Fig.3E). TH immunohistochemistry and TH immunofluorescence staining (Fig.3F,3G, 3H, 3I) revealed that in the model group, the number of TH-positive-neurons in the lesioned substantia nigra was significantly reduced, and the TH fluorescence signal was markedly weakened. Following TH⁺/FOXA2⁺ cell-derived EVs treatment, the number of TH-positive-neurons significantly increased, and the TH fluorescence intensity was notably restored. Notably, the extent of TH restoration did not increase monotonically with dose: in the quantitative analyses shown in Fig. 3H and 3I, the high-dose group tended to show a smaller increment in TH-positive neuron number and TH fluorescence intensity than the medium-dose group. A plausible explanation is that TH⁺/FOXA2⁺ cell-derived EVs are of human origin, so that a higher cumulative burden of xenogeneic human-derived EVs is more likely to elicit a host immune response in the recipient animals — a tendency that is also suggested by the dose-related attenuation observed in A53T mice. Such an immune response (for example, microglial activation, anti-EV antibody production and complement-mediated opsonization) would accelerate the clearance of the administered EVs from the brain, shorten their effective residence time and thereby partially offset the reparative effect. This interpretation implies that the dose–response relationship of TH⁺/FOXA2⁺ cell-derived EVs is not linear and that an optimal dose window, rather than the maximum achievable dose, needs to be defined. Direct measurement of anti-EV antibodies, complement activation and neuroinflammatory markers across dose levels will be required to confirm this hypothesis. Collectively, these results demonstrate that TH⁺/FOXA2⁺ cell-derived EVs effectively ameliorate motor dysfunction and alleviate dopaminergic neuron injury in the substantia nigra of 6-OHDA-induced--PD rats.

#### 3.2.3 TH⁺/FOXA2⁺ cell-derived EVs improve brain structural abnormalities in 6-OHDA-induced PD rats

Following histological assessment, we further validated the therapeutic effects of TH⁺/FOXA2⁺ cell-derived EVs at the levels of whole-brain-structure and metabolic function using magnetic resonance imaging (MRI) and positron emission tomography/computed-tomography (PET/CT-). MRI results showed that the ventricular volume in the model group was significantly increased compared with the control group, indicating brain atrophy or tissue loss. Compared with the model group, the ventricular volume was markedly improved in the TH⁺/FOXA2⁺ cell-derived EVs-HD- and Madopar groups; although the TH⁺/FOXA2⁺ cell-derived EVs-MD- and EVs-LD groups also exhibited a decreasing trend in ventricular volume, the difference was not statistically significant compared with the model group, suggesting that high—dose TH⁺/FOXA2⁺ cell-derived EVs and Madopar effectively attenuated ventriculomegaly and ameliorated structural brain damage in the PD rat model (Fig. 4A, 4C). PE-T/CT results showed that the model group exhibited a decreasing trend in the striatum-to-cerebe--llum standardized uptake value ratio (SUVr) compared with the control group, indicating reduced dopamine synthesis activity, which is consistent with the expected direction of decreased dopamine-rel-ated tracer signals following 6--OHDA lesion. Compared with the model group, the TH⁺/FOXA2⁺ cell-derived EVs dose groups and the Madopar positive control group showed no statistically significant differences, but all exhibited varying degrees of increasing trends, suggesting a certain tendency toward improvement in brain imaging signals following intervention (Fig. 4B, 4D).

**Figure 4.**
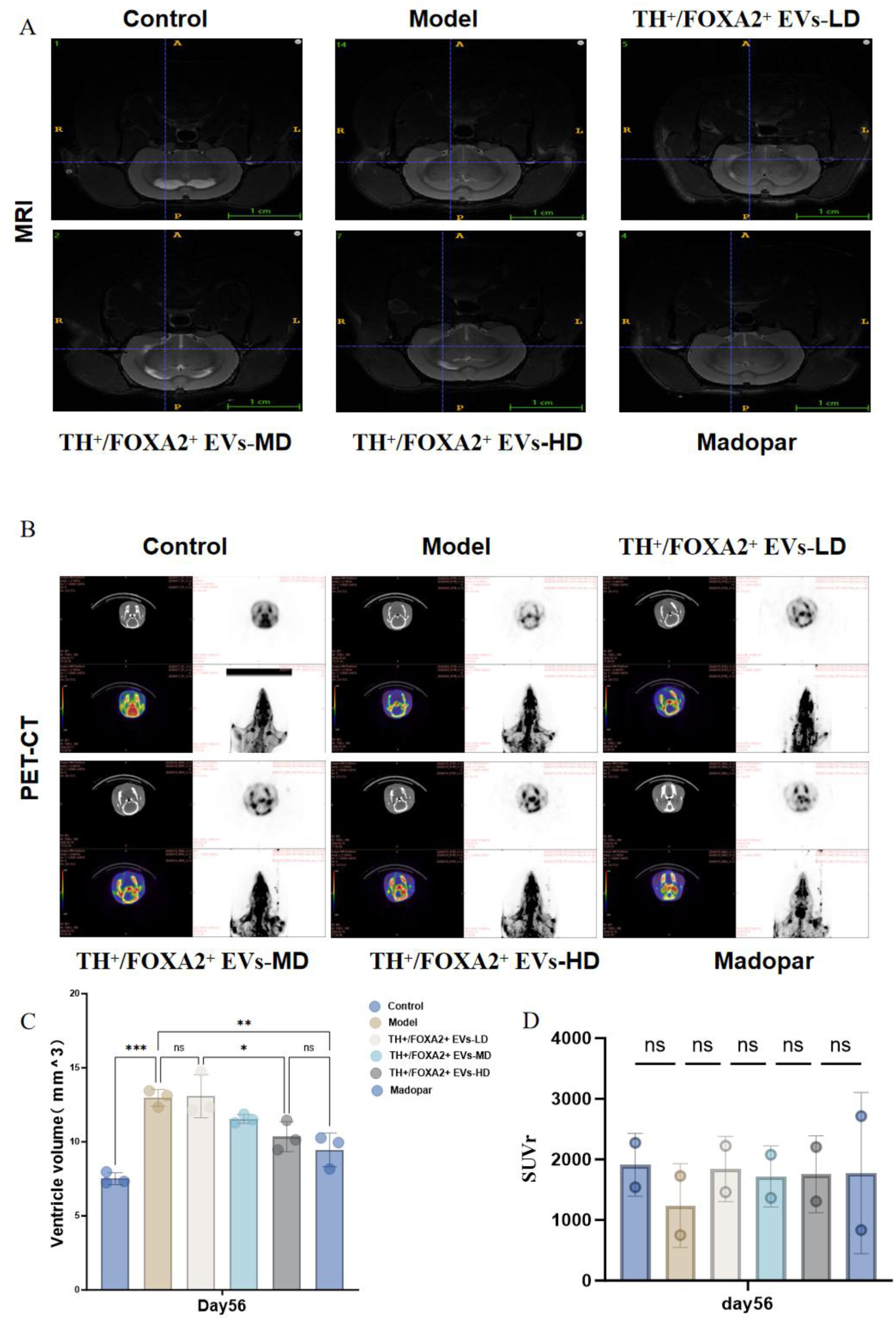
TH⁺/FOXA2⁺ cell-derived EVs improve brain structural abnormalities in 6-OHDA-induced PD rats. TH⁺/FOXA2⁺ cell-derived EVs improve brain structural abnormalities in 6-OHDA-induced--PD rats. (A) MRI results of PD rats treated with different doses of TH⁺/FOXA2⁺ cell-derived EVs for 8 weeks. (B) PET/CT-results of PD rats treated with different doses of TH⁺/FOXA2⁺ cell-derived EVs for 8 weeks. (C) Quantitative analysis of MRI in PD rats treated with different doses of TH⁺/FOXA2⁺ cell-derived EVs for 8 weeks (n = 3). (D) Quantitative analysis of PET/CT- in PD rats treated with different doses of TH⁺/FOXA2⁺ cell-derived EVs for 8 weeks (n = 2). Scale bar = 1 cm. Data are shown as mean ± SD. ns, not significant; *p<0.05, **p<0.01, ***p<0.001 vs. model group. (A) and (B) are representative images from n = 3 (MRI) and n = 2 (PET/CT) biologically independent animals per group. All n values refer to biologically independent animals (biological replicates).

### 3.3 In Vitro Experiments

#### 3.3.1 TH⁺/FOXA2⁺ cell-derived EVs Ameliorate Dopaminergic Neuronal Damage in the 6-OHDA-Induced MBO PD Model--

Before PD modeling, the identity of the MBOs was verified by immunofluorescence. Organoids at day 60 showed strong and regionally organized expression of the floor-plate/midbrain markers FOXA2, LMX1A and CORIN, together with the dopaminergic markers TH, NURR1 and DAT and the pan-neuronal marker TUJ1, while the pluripotency marker OCT4 was essentially undetectable, confirming the successful generation of ventral midbrain-floor-plate-patterned organoids that contain dopaminergic neurons.

To verify the effect of TH⁺/FOXA2⁺ cell-derived EVs *in vitro* and across species differences, we established an *in vitro* PD model using 6-OHDA-treated--human midbrain organoids (MBOs). On day 60 of MBO culture, the PD model was generated by co-culturing-MBOs with 300μM 6-OHDA-for 48 h (Fig.5A,5B). Subsequently, two intervention strategies were compared: co-culture-with 5×10¹⁰ particles/mL of TH⁺/FOXA2⁺ cell-derived EVs during PD modeling (TH⁺/FOXA2⁺ cell-derived EVs-2-), or addition of the same concentration of TH⁺/FOXA2⁺ cell-derived EVs after modeling (TH⁺/FOXA2⁺ cell-derived EVs-1-) (Fig.5B). Thus, in the TH⁺/FOXA2⁺ cell-derived EVs-1 group, EV treatment was initiated only after the completion of 6-OHDA modeling (post-modeling treatment), whereas in the TH⁺/FOXA2⁺ cell-derived EVs-2 group, EVs were administered at the same time as the modeling insult, i.e., modeling and treatment were performed simultaneously. As shown below, this simultaneous modeling-and-treatment regimen (TH⁺/FOXA2⁺ cell-derived EVs-2) produced a stronger therapeutic effect than the post-modeling regimen (TH⁺/FOXA2⁺ cell-derived EVs-1). After 48 h, both groups continued to receive 5×10¹⁰ particles/mL of TH⁺/FOXA2⁺ cell-derived EVs for an additional 7 days. Following modeling, MAP2-positive-neurons exhibited sparse and fragmented dendritic structures, and the number of TH-positive-dopaminergic neurons was significantly reduced. After treatment with TH⁺/FOXA2⁺ cell-derived EVs, the integrity of the MAP2-positive-dendritic network was markedly restored, and the number of TH-positive-neurons was also significantly increased in the TH⁺/FOXA2⁺ cell-derived EVs-2-group, indicating that TH⁺/FOXA2⁺ cell-derived EVs protect the morphological structure of dopaminergic neurons (Fig.5C, 5D, 5E).

**Figure 5.**
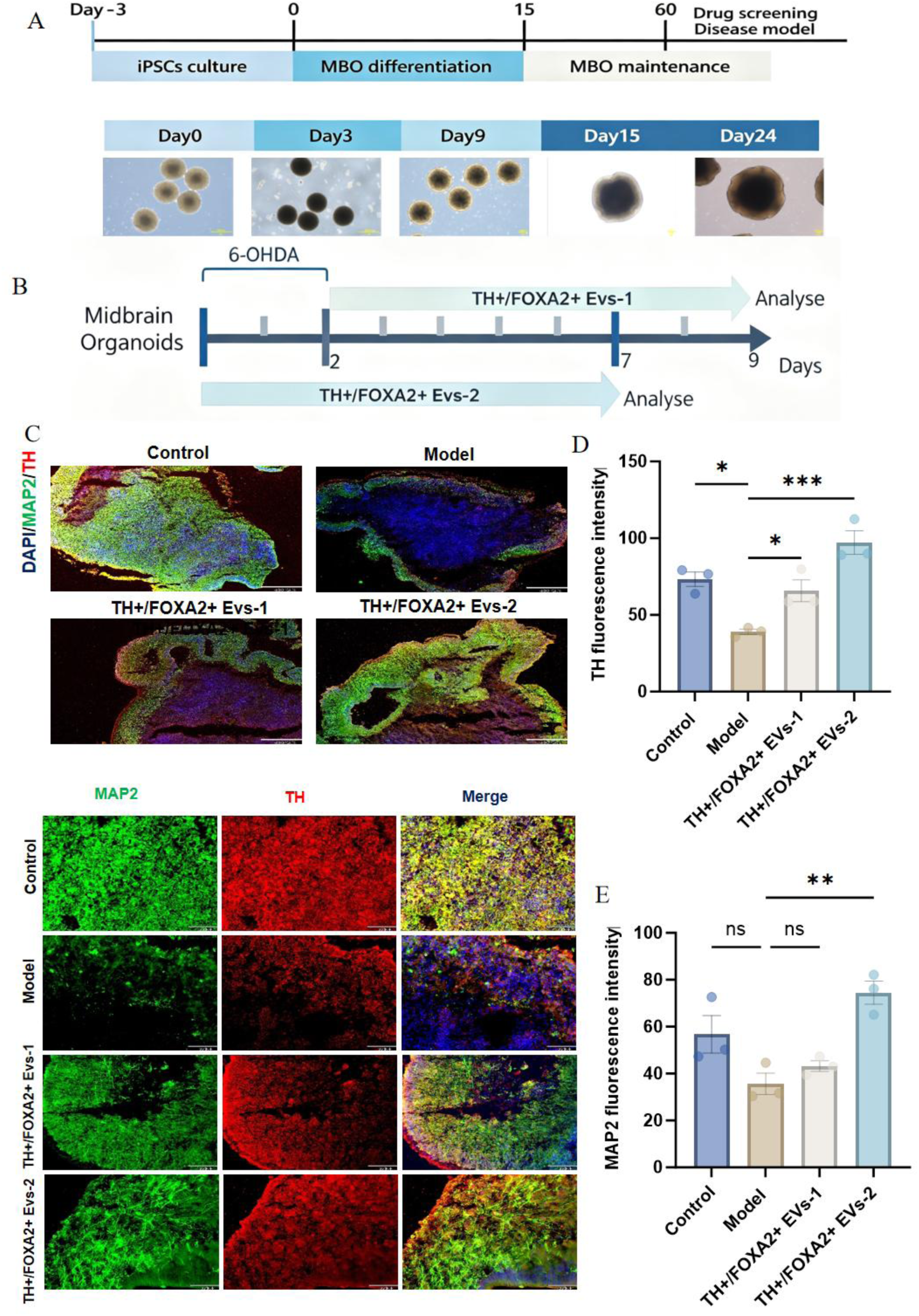
TH⁺/FOXA2⁺ cell-derived EVs ameliorate dopaminergic neuronal damage in the midbrain organoid PD model. TH⁺/FOXA2⁺ cell-derived EVs ameliorate neuronal damage in the MBO PD model. (A) Workflow for establishing the MBO PD model. (B) Different protocols for 6-OHDA-modeling. (C) MAP2/TH immunofluorescence staining of MBOs from control, model, and TH⁺/FOXA2⁺ cell-derived EVs-treated-groups with different intervention durations. (D) Quantitative analysis of MAP2 staining in MBOs from control, model, and TH⁺/FOXA2⁺ cell-derived EVs-treated-groups with different intervention durations (n = 3). (E) Quantitative analysis of TH staining in MBOs from control, model, and TH⁺/FOXA2⁺ cell-derived EVs-treated-groups with different intervention durations (n = 3). Scale bar = 90μm. ns, not significant; *p<0.05, **p<0.01, ***p<0.001. (C) is a representative image from n = 3 biologically independent organoid batches; all quantitative panels show n = 3 biological replicates. Groups: control (no 6-OHDA, no EVs); model (6-OHDA only); TH⁺/FOXA2⁺ cell-derived EVs-1 (post-modeling treatment, EVs given only after modeling was completed); TH⁺/FOXA2⁺ cell-derived EVs-2 (concurrent treatment, EVs given simultaneously with 6-OHDA modeling). After the 48-h modeling period, both EV groups received EVs for a further 7 days. Among them, the simultaneous modeling-and-treatment group (TH⁺/FOXA2⁺ cell-derived EVs-2) showed the stronger protective effect.

#### 3.3.2 TH⁺/FOXA2⁺ cell-derived EVs improve glial-neuronal colocalization in the 6-OHDA-induced MBO PD model

To further evaluate the effects of TH⁺/FOXA2⁺ cell-derived EVs, GFAP/TUJ1 double staining and Western blotting were performed to assess the impact of TH⁺/FOXA2⁺ cell-derived EVs-1-and TH⁺/FOXA2⁺ cell-derived EVs-2-on glial activation and neuronal damage. GFAP/TUJ1 co-staining-showed that in the model group, GFAP-positive-astrocytes were significantly activated, the number of TUJ1-positive-neurons was reduced, and their co-localization was abnormal (Fig.6A,6B,6C). Both TH⁺/FOXA2⁺ cell-derived EVs-1-and TH⁺/FOXA2⁺ cell-derived EVs-2-treatments significantly decreased GFAP fluorescence intensity and increased the number of TUJ1-positive-neurons. Western blotting results showed that TH protein levels were significantly reduced in the model group, and both TH⁺/FOXA2⁺ cell-derived EVs-1-and TH⁺/FOXA2⁺ cell-derived EVs-2-markedly restored TH expression, with comparable efficacy between the two treatments (Fig.6D,6G). GFAP protein levels were elevated in the model group and were significantly downregulated only by TH⁺/FOXA2⁺ cell-derived EVs-2-, while the TH⁺/FOXA2⁺ cell-derived EVs-1-group showed no statistically significant difference from the model group (Fig. 6D, 6E). Similarly, α-syn-aggregation was increased in the model group, and only TH⁺/FOXA2⁺ cell-derived EVs-2-significantly reduced its expression, with no significant effect observed for TH⁺/FOXA2⁺ cell-derived EVs-1-(Fig. 6D, 6F). In summary, both TH⁺/FOXA2⁺ cell-derived EVs-1-and TH⁺/FOXA2⁺ cell-derived EVs-2-effectively restored TH expression and improved glial-neuronal-co-localization; however, TH⁺/FOXA2⁺ cell-derived EVs-2-exhibited stronger anti-gliotic-and anti-α-syn--aggregation capabilities in terms of suppressing GFAP and α-syn-protein levels.

**Figure 6.**
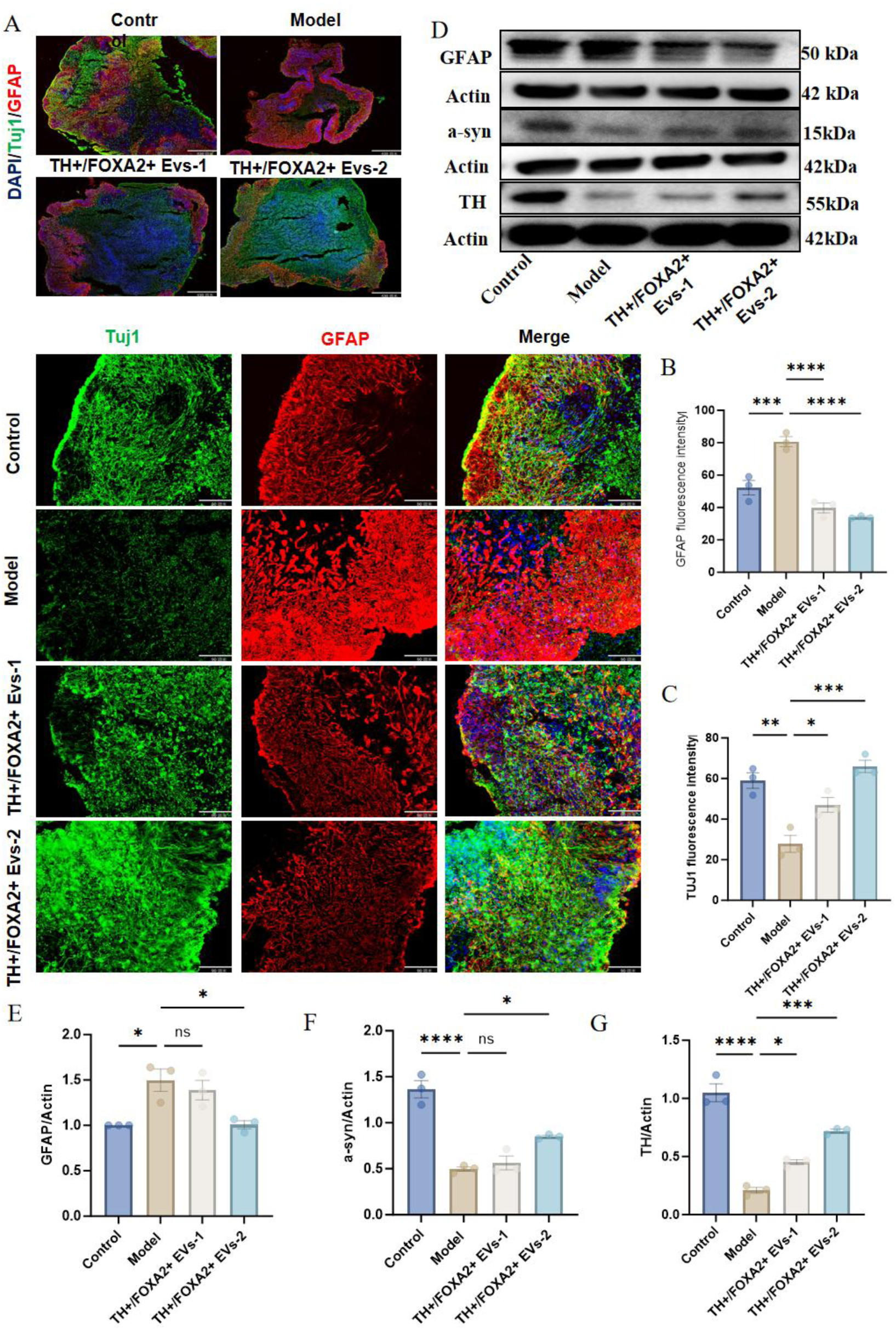
TH⁺/FOXA2⁺ cell-derived EVs improve glial-neuronal colocalization in the 6-OHDA-induced MBO PD model. TH⁺/FOXA2⁺ cell-derived EVs improve glial-neuronal-colocalization in the 6-OHDA-induced--MBO PD model. (A) TUJ1/GFAP immunofluorescence staining of MBOs from control, model, and TH⁺/FOXA2⁺ cell-derived EVs-treated-groups with different intervention durations. (B) Quantitative analysis of GFAP staining in MBOs from control, model, and TH⁺/FOXA2⁺ cell-derived EVs-treated-groups with different intervention durations (n = 3). (C) Quantitative analysis of TUJ1 staining in MBOs from control, model, and TH⁺/FOXA2⁺ cell-derived EVs-treated-groups with different intervention durations (n = 3). (D) Western blot analysis of protein expression in MBOs from control, model, and TH⁺/FOXA2⁺ cell-derived EVs-treated-groups with different intervention durations. (E) Quantification of GFAP protein expression in MBOs from control, model, and TH⁺/FOXA2⁺ cell-derived EVs-treated-groups with different intervention durations (n = 3). (F) Quantification of α-syn-protein expression in MBOs from control, model, and TH⁺/FOXA2⁺ cell-derived EVs-treated-groups with different intervention durations (n = 3). (G) Quantification of TH protein expression in MBOs from control, model, and TH⁺/FOXA2⁺ cell-derived EVs-treated-groups with different intervention durations (n = 3). Scale bar = 90 μm. ns, not significant; *p<0.05, **p<0.01, ***p<0.001. (A) and (D) are representative images from n = 3 biologically independent organoid batches; all quantitative panels show n = 3 biological replicates. Groups: control (no 6-OHDA, no EVs); model (6-OHDA only); TH⁺/FOXA2⁺ cell-derived EVs-1 (post-modeling treatment, EVs given only after modeling was completed); TH⁺/FOXA2⁺ cell-derived EVs-2 (concurrent treatment, EVs given simultaneously with 6-OHDA modeling). Overall, the TH⁺/FOXA2⁺ cell-derived EVs-2 group, in which modeling and treatment were performed simultaneously, exhibited stronger efficacy in suppressing glial activation and α-syn accumulation than the TH⁺/FOXA2⁺ cell-derived EVs-1 post-modeling treatment group.

## 4. Multi-Omics and Single-Cell Sequencing Results of TH⁺/FOXA2⁺ cell-derived EVs Treatment in the 6-OHDA-Induced MBO PD Model---

To elucidate the molecular mechanisms of TH⁺/FOXA2⁺ cell-derived EVs, transcriptomic and single-cell-sequencing were performed on MBOs from the control, model, TH⁺/FOXA2⁺ cell-derived EVs-1-, and TH⁺/FOXA2⁺ cell-derived EVs-2-groups. Transcriptomic sequencing results revealed that a total of 116 differentially expressed genes were identified in the TH⁺/FOXA2⁺ cell-derived EVs-2-treatment group compared with the control and model groups. We focused on genes associated with neuroinflammation and neuronal/synaptic function. Regarding inflammation-related-genes, the TH⁺/FOXA2⁺ cell-derived EVs-2-treatment group significantly regulated multiple key immune modulators, including ferritin light chain (FTL), glial fibrillary acidic protein (GFAP), beta-2-microglobulin-(B2M), interferon-induced-transmembrane protein 3 (IFITM3), interferon-stimulated-gene 15 (ISG15), human leukocyte antigens HLA-C- and HLA-B-, secreted phosphoprotein 1 (SPP1), chemokine ligand 2 (CCL2), complement C1q-like-protein 1 (C1QL1), interferon regulatory factor 7 (IRF7) and IRF9, as well as galectin-3- (LGALS3). Concurrently, multiple genes associated with neuronal structure and synaptic function were also significantly altered, including synaptosome-associated protein 25 (SNAP25), neurofilament -medium chain (NEFM), contactin 4 (CNTN4), neurexin 1 (NRXN1), glutamate decarboxylase 1 (GAD1), synaptotagmin 4 (SYT4), RNA binding motif protein 25 (RBM25), GA-binding protein transcription fa-ctor α subunit (GABPA), embryonic lethal abnormal vision-like protein 4 (ELAVL4), cont-actin-associated protein 2 (CNTNAP2), vis-inin-like protein 1 (VSNL1), mitog-en-activated protein kinase 3K19 (MAP-3K19), zinc finger protein 638 (ZNF638), and TRAF family member-associated NF-κB activator (TANK). -These -alterations in gene expression suggest that TH⁺/FOXA2⁺ cell-derived EVs-2 may exert therapeutic effects on 6-OHDA-induced P-D organoids through coordinated regulation of ge--nes involved in inflammatory responses and neuronal function.

To validate the establishment of the MBO PD model, we performed single-cell-sequencing on the organoids. A total of 24 cell subpopulations were annotated, among which astrocytes (AQP4, GFAP, GJA1, SLC1A2, SLC1A3) accounted for approximately 38.2%, floor plate cells (FOXA2, SHH, CORIN, LMX1B) accounted for 14.2%, and neurons (INA, SYT1, SNAP25) accounted for 7.6%, fully recapitulating the developmental lineage of midbrain dopaminergic neurons from floor plate precursors to mature neurons (Fig.7A). Subsequently, each subpopulation was annotated with specific marker genes. The results showed that the core functional cells of these organoids are dopaminergic (DA) neurons, together with their development-related-precursor cells, neurogenic cells, and cells derived from the ventral midbrain floor plate (FP). Marker genes of mature DA neurons (TH, DDC, DAT) were specifically and highly expressed in the DA neuron cluster; marker genes of the midbrain floor plate and neural precursors (FOXA2, LMX1B, DCX) were enriched in the corresponding developmental stage cell populations. This overall expression pattern fully recapitulates the lineage progression of midbrain DA neurons from development to maturation, consistent with the biological characteristics of midbrain organoids (Fig.7B).

**Figure 7.**
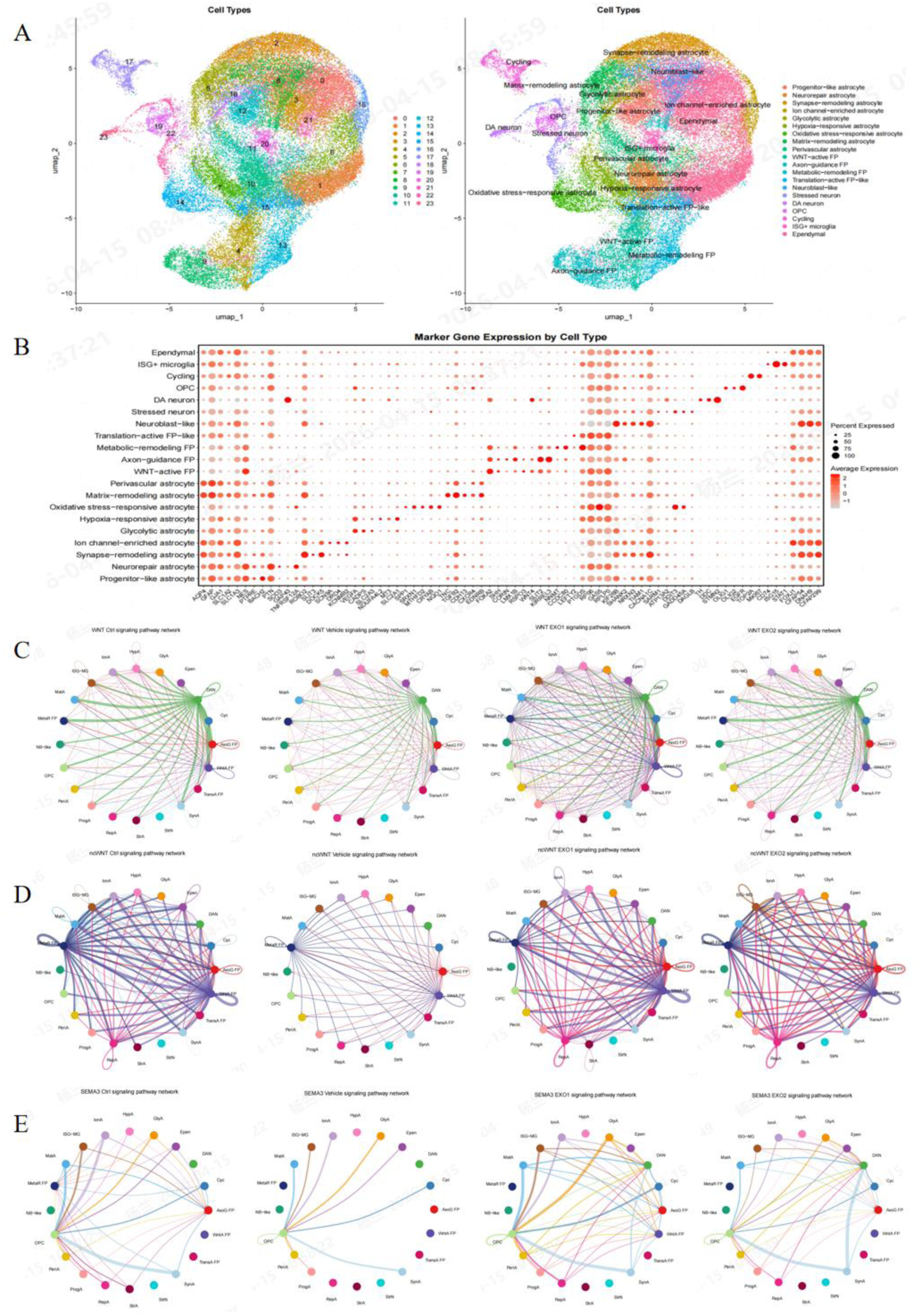
Protective Mechanism of TH⁺/FOXA2⁺ cell-derived EVs for Dopaminergic Neurons in Parkinson’s Disease Model Organoids Based on Single-Cell Sequencing. Single-cell-sequencing results of TH⁺/FOXA2⁺ cell-derived EVs treatment in the 6-OHDA-induced--MBO PD model. (A) Cell annotation of the MBO PD model. (B) Marker gene annotation of cell subpopulations in the MBO PD model. (C) Cell--cell communication network of the WNT pathway in control, model, and TH⁺/FOXA2⁺ cell-derived EVs-treate-d MBO PD models with different intervention durations. (D) Cel-l-cell communication network of the ncWNT pathway. (E) Ce-ll-cell communication network of the SEMA3 pathway. n = 3 biologically independent organoid samples per group (biological replicates).

Further, we performed cell-cell-communication analysis. The WNT pathway is a key regulatory pathway for midbrain DA neuron differentiation and floor plate specification. In the PD model, intercellular communication was significantly weakened; however, after TH⁺/FOXA2⁺ cell-derived EVs treatment, the intercellular connections became significantly more numerous and denser, particularly manifested as enhanced interactions among DA neurons, astrocytes (Astro), floor plate cells (FP), and vesicular monoamine transporter-positive-floor plate cells (Vmat⁺ FP). This suggests that TH⁺/FOXA2⁺ cell-derived EVs may activate WNT signaling-mediated-cell communication, thereby promoting the differentiation of midbrain floor plate precursor cells into DA neurons or enhancing the supportive role of glial cells toward neurons (Fig.7C). Regarding the non-canonical-WNT (ncWNT) pathway, this pathway is generally involved in nutritional support and injury responses among vascular endothelial cells, astrocytes, and neurons. Compared with the model group, TH⁺/FOXA2⁺ cell-derived EVs treatment reversed the inhibition of the ncWNT pathway, restored the normal function of astrocytes, enhanced their nutritional support and homeostatic regulation communication with neurons and other cells, and improved the post-injury microenvironment (Fig.7D). The SEMA3 pathway primarily regulates axonal growth, migration, and synapse formation of neurons and is critical for the projection and maturation of DA neurons. Compared with the control group, the overall communication intensity of the SEMA3 pathway was reduced in the model group, with some intercellular connections disappearing; the most pronounced reduction was observed in the interaction between oligodendrocyte precursor cells (OPCs) and DA neurons, suggesting that axonal guidance and myelination-related-communication were inhibited in the model group, potentially leading to impaired neuronal maturation. After TH⁺/FOXA2⁺ cell-derived EVs treatment, the cell communication of the SEMA3 pathway was restored and enhanced, likely promoting axonal growth, migration, and myelination of DA neurons and improving the maturation status of neurons (Fig.7E).

Single-cell-transcriptomics combined with transcriptomic analysis further revealed the molecular signatures of specific glial subpopulations. In interferon-responsive-microglia (ISG⁺ microglia), analysis of the top 20 differentially expressed genes (DEGs) showed that IFITM3, ISG15, IFI6, and STAT1 were significantly upregulated, suggesting that they may serve as key stress-responsive-factors. Meanwhile, among the top 20 DEGs in oxidative stress-adapted astrocytes, FTL, CRYAB, -EIF4EBP1, CDKN1A, and SPP1 were aberrantly expressed, potentially constituting pathogenic or stress-related molecules.-

## 5. Toxicological Evaluation of TH⁺/FOXA2⁺ cell-derived EVs Treatment in PD Models

During the study period, no death or moribund status was observed in rats or cynomolgus monkeys. Clinical observations (including general clinical observation, detailed clinical observation, local observation at the administration site), body weight, body temperature, blood pressure, hematology, coagulation function, serum biochemistry, urinalysis, C-reactive-protein, immunophenotyping, and immunoglobulins, as well as gross necropsy findings in rats and cynomolgus monkeys, revealed no abnormal changes related to TH⁺/FOXA2⁺ cell-derived EVs.

## 6. Discussion

In recent years, significant progress has been made in the use of extracellular vesicles (EVs) for the treatment of Parkinson’s disease, encompassing engineered EVs, mesenchymal stem cell-derived-EVs (MSC-EVs), and the use of EVs as biomarkers[19–24]. In comparison, TH⁺/FOXA2⁺ cell-derived EVs are derived from midbrain floor-plate progenitors differentiating into a population enriched with DA neurons and region-specific glial cells, thereby endowing TH⁺/FOXA2⁺ cell-derived EVs with more “tissue-matched-” repair signals. To substantiate this, we performed transcriptomic and proteomic analyses of these EVs and conducted an in-depth-comparison with conventional MSC-EVs. We found that TH⁺/FOXA2⁺ cell-derived EVs are enriched in a series of molecules critical for neuronal function, survival, and synaptic plasticity. For example, STMN2, a key regulator of neuronal plasticity and axonal regeneration, is downregulated in neurons of Parkinson’s disease patients, leading to impaired axonal repair[25, 26]. Delivery of STMN2 by TH⁺/FOXA2⁺ cell-derived EVs may directly repair damaged axonal transport and the cytoskeleton, which explains the improved motor function observed in our *in vivo* models. SPARCL1 is a known astrocyte-derived synaptogenic fa-ctor[27], and BCAN is a core component of the perineuronal net that maintains synaptic function by stabilizing synaptic structure and modulating AMPA receptor activity[28]. Their high expression in TH⁺/FOXA2⁺ cell-derived EVs suggests that these EVs may participate in synaptic maintenance and formation, which is critical for the restoration of the lost synaptic connections in PD. The therapeutic efficacy of TH⁺/FOXA2⁺ cell-derived EVs is not attributable to a single molecule but rather results from the coordinated action of proteins, mRNAs, and miRNAs, simultaneously targeting axonal regeneration, synaptic maintenance, cytoskeletal stability, and anti-inflammation, thereby conferring a -multidimensional therapeutic advantage that warrants comparative validation in future studies.

An important conceptual point should be emphasized regarding the cellular source of TH⁺/FOXA2⁺ cell-derived EVs. The TH⁺/FOXA2⁺ cells used for EV production are not conventional, terminally differentiated midbrain dopaminergic (A9) neurons. They are generated from human iPSCs through a midbrain floor-plate differentiation route and represent a distinct cell population that co-expresses the dopaminergic marker TH together with the floor-plate and ventral midbrain progenitor transcription factor FOXA2. Persistent FOXA2 expression indicates retention of floor-plate/progenitor identity, whereas bona fide mature substantia nigra dopaminergic neurons are defined by the coordinated expression of TH, AADC (DDC), DAT (SLC6A3), NURR1 (NR4A2), PITX3 and GIRK2 (KCNJ6), together with extinction of the FOXA2/LMX1A progenitor program. The TH⁺/FOXA2⁺ population used here is therefore best described as a TH⁺/FOXA2⁺ midbrain floor-plate-derived neural cell population at an intermediate differentiation stage rather than as classical mature dopaminergic neurons. Accordingly, the cargo of TH⁺/FOXA2⁺ cell-derived EVs reflects the developmental and reparative secretory program of this specific population, rather than the secretome of adult dopamine neurons, which provides the mechanistic basis for the trophic, pro-regenerative and anti-inflammatory effects observed in the present study.

This study systematically evaluated, for the first time, the application potential of TH⁺/FOXA2⁺ cell-derived EVs in the treatment of Parkinson’s disease. As a potential biologic agent, the safety of TH⁺/FOXA2⁺ cell-derived EVs is a prerequisite for their clinical translation. Our toxicological results demonstrated that at therapeutically effective doses, TH⁺/FOXA2⁺ cell-derived EVs did not induce obvious hepatorenal toxicity (normal levels of ALT, AST, BUN, and Cr), nor did they trigger significant acute inflammatory responses or pathological damage in major organs. These findings preliminarily confirm the favorable safety profile of TH⁺/FOXA2⁺ cell-derived EVs, laying a foundation for their subsequent clinical development. In animal experiments, intranasal administration of TH⁺/FOXA2⁺ cell-derived EVs significantly improved motor coordination and rotational behavior, and increased the number of TH-positive-neurons in the substantia nigra. Notably, MRI showed a trend toward reduced ventricular volume in the medium-dose-group, with a significant reduction achieved by the high dose. Furthermore, comparison of the onset of efficacy between the two models revealed that in A53T mice, different dose groups of TH⁺/FOXA2⁺ cell-derived EVs exhibited sustained effects starting from day 7, whereas in 6-OHDA-rats, consistent efficacy across the behavioral endpoints was achieved from day 14 onward. This difference may stem from the distinct pathological mechanisms: the A53T model is -characterized by chronic α-synuclein aggregation with a relatively slow neurodegenerative-progression[29], whereas the 6-OHDA model involves acute oxidative stress injury[30], requiring a longer period for EVs to establish a reparative microenvironment. This suggests that TH⁺/FOXA2⁺ cell-derived EVs may exert superior intervention effects on chronic, progressive pathological processes compared to acute injury-. In the MBO model, TH⁺/FOXA2⁺ cell-derived EVs-2 was significantly superior to TH⁺/FOXA2⁺ EVs-1 in suppressing GFAP and α-syn. This finding indicates that the therapeutic efficacy of TH⁺/FOXA2⁺ cell-derived EVs is highly dependent on the timing of intervention, implying that their clinical application may need to target prodromal PD patients. Specifically, administering TH⁺/FOXA2⁺ cell-derived EVs simultaneously with 6-OHDA exposure (TH⁺/FOXA2⁺ cell-derived EVs-2), rather than initiating treatment only after the modeling insult had been completed (TH⁺/FOXA2⁺ cell-derived EVs-1), produced markedly stronger protection. This indicates that performing EV treatment at the same time as the insult confers better efficacy than delayed intervention after modeling, most likely because EVs are present while the injury cascade is being initiated, allowing early suppression of oxidative damage, glial activation and pathological protein deposition before irreversible neuronal loss occurs.

From a translational perspective, a further advantage of the present platform resides in its manufacturing process. Conventional protocols for generating midbrain dopaminergic cells or brain organoids generally rely on extracellular matrix substrates—such as Matrigel or Geltrex—for cell attachment or embedding, and depend on proprietary supplements, most notably N2 and B27, as indispensable components of the neural induction and maturation media. These reagents markedly increase the cost of goods, introduce undefined components of animal origin, and exhibit appreciable lot-to-lot variability, which together complicate process scale-up and compromise batch-to-batch consistency. In contrast, the differentiation system established here is suspension-based throughout: from EB formation on day 0 until harvest, the cells are maintained as free-floating aggregates in ultra-low attachment vessels in a chemically defined medium that contains neither N2 nor B27 and requires no matrix coating. This design simplifies the workflow, minimizes manual handling and operator-dependent variation, lowers reagent cost, and is readily amenable to further scale-up. Together with the tangential flow filtration-based purification described above, it delivers a simple, controllable, low-cost and scalable production route with high batch-to-batch consistency, a critical quality attribute for the clinical translation of EV-based biologics.

To bridge the species gap between rodent models and human disease, and to establish an experimental system that more closely recapitulates the complex cellular composition and three-dimensional-architecture of the human brain, we developed a 6-OHDA-induced--human MBO PD model. GFAP, a classic marker of astrocyte activation, was highly expressed, directly reflecting reactive gliosis and the remodeling of the neuroinflammatory microenvironment under PD pathological conditions. This finding is particularly important because neuroinflammation is not merely a downstream product of PD pathology but also a key driver of disease progression; activated astrocytes release inflammatory factors that exacerbate dopaminergic neuronal damage[31–33]. Treatment with 300μM 6-OHDA-for 48 h successfully induced neuronal damage (reduced MAP2/TH-positive-structures), and intervention with TH⁺/FOXA2⁺ cell-derived EVs effectively restored TH expression and ameliorated the abnormal GFAP/TUJ1 co-localization, indicating that TH⁺/FOXA2⁺ cell-derived EVs can restore the survival and morphology of dopaminergic neurons. Interestingly, when TH⁺/FOXA2⁺ cell-derived EVs were added at the same time as modeling (TH⁺/FOXA2⁺ cell-derived EVs-2-group) rather than after modeling (TH⁺/FOXA2⁺ cell-derived EVs-1-group), their inhibitory effects on GFAP protein and α-syn-aggregation were more pronounced, suggesting that early intervention may be more favorable for suppressing glial activation and pathological protein deposition[34–36]. This indicates that the therapeutic effects of TH⁺/FOXA2⁺ cell-derived EVs are not limited to direct neuronal protection but also profoundly influence the functional state of glial cells. These findings provide the first evidence for the protective effects of TH⁺/FOXA2⁺ cell-derived EVs in a complex three-dimensional-human neural tissue model, not only validating the observations from animal experiments but also offering human-specific-molecular-level-evidence, thereby greatly enhancing the clinical relevance of the study.

To further elucidate the molecular basis of the specific therapeutic effects of TH⁺/FOXA2⁺ cell-derived EVs, we performed transcriptomic and single-cell-sequencing analyses on the MBOs and identified significant upregulation of IFITM3, ISG15, SNAP25, NEFM, and NRXN1. IFITM3 and ISG15 are canonical downstream effector molecules of the type I interferon (IFN-I-) signaling pathway. Studies have shown that IFN-I-signaling is aberrantly activated in the brains of PD patients and in cells with α-synuclein-aggregation, and this activation is closely associated with persistent microglial activation in the substantia nigra[37, 38]. TH⁺/FOXA2⁺ cell-derived EVs significantly reduced the expression of IFITM3/ISG15, suggesting that they may alleviate neuroinflammation through negative regulation of the IFN-I-pathway. Among the differentially expressed genes related to neuronal function in our analysis, SNAP25, NEFM, and NRXN1 were significantly upregulated. SNAP25 is a core component of the presynaptic SNARE complex, and its decreased expression directly leads to impaired dopamine release[39, 40]; NEFM is the neurofilament medium chain that maintains axonal stability[41, 42]; and NRXN1 is involved in synaptic specification and maintenance[43, 44]. The restoration of these genes provides direct molecular evidence for the synaptic protection observed in our behavioral and TH immunohistochemical analyses.

In the single-cell-sequencing analysis, this study focused on three key intercellular communication pathways. In the PD model group, the communication intensity of the WNT pathway was significantly weakened, with the connection between floor plate cells and dopaminergic neurons almost disappearing. WNT/β-catenin-signaling is essential for midbrain floor plate specification and terminal differentiation of DA neurons[45, 46]. After TH⁺/FOXA2⁺ cell-derived EVs treatment, WNT communication between FP and DA neurons was markedly enhanced. Concurrently, we observed that the non-canonical-WNT (ncWNT) pathway, which primarily mediates metabolic support from astrocytes to neurons[47], also recovered its activity. Thus, TH⁺/FOXA2⁺ cell-derived EVs may partially reprogram mature glial cells to promote neuronal repair by reactivating developmental WNT signaling. Members of the SEMA3 family regulate axonal growth cone guidance and presynaptic differentiation by binding to neuropilin receptors. In the PD model, overall communication via the SEMA3 pathway was diminished, with the most pronounced reduction observed in the connection between oligodendrocyte precursor cells (OPCs) and DA neurons. OPC-derived-SEMA3 signals can restrict aberrant axonal sprouting and maintain synaptic specificity[48]. Following TH⁺/FOXA2⁺ cell-derived EVs treatment, this pathway was significantly restored, consistent with the observed increases in TH-positive-fiber density and MAP2 dendritic integrity.

Integration of transcriptomic and single-cell-sequencing analyses further annotated a distinct population of interferon-responsive-microglia. The expression levels of IFITM3, ISG15, IFI6, and STAT1 were elevated in the PD model group but were significantly reduced after TH⁺/FOXA2⁺ cell-derived EVs treatment. This is highly consistent with the “disease-associated-microglia” or “IFN-responsive-glia” recently reported in multiple single-cell-studies of PD patient brains[49]. TH⁺/FOXA2⁺ cell-derived EVs may block the transmission of inflammation to neurons by suppressing the expansion of this subpopulation or reducing its IFN signaling intensity. In addition, we identified a population of oxidative stress-adapted-astrocytes characterized by high expression of FTL, CRYAB, EIF4EBP1, CDKN1A, and SPP1. CRYAB is a small heat shock protein that protects the cytoskeleton under oxidative stress[50]; CDKN1A (p21) is a cell cycle inhibitor that can induce a senescence-associated-secretory phenotype (SASP)[51]. After TH⁺/FOXA2⁺ cell-derived EVs treatment, the aberrant expression profile of this subpopulation tended to normalize, suggesting that the treatment alleviated stress-induced-senescence of astrocytes, thereby improving glial-neuronal-metabolic coupling. We propose that TH⁺/FOXA2⁺ cell-derived EVs may first suppress the IFN-I-signaling of interferon-responsive-microglia, subsequently alleviating the oxidative stress/senescence phenotype of astrocytes. The reshaped glial microenvironment then creates favorable conditions for the restoration of WNT and SEMA3 signaling pathways, ultimately promoting DA neuron survival, axonal growth, and synaptic maintenance.

Despite the substantial potential of TH⁺/FOXA2⁺ cell-derived EVs demonstrated in PD models, the present study has the following limitations. TH⁺/FOXA2⁺ cell-derived EVs are derived from human midbrain floor-plate progenitors but were administered in rodents. Although clear therapeutic effects were observed, species differences in surface proteins of EVs may affect their targeting efficiency and bioavailability, and the possibility of xenogeneic immune reactions cannot be completely ruled out. Secondly, the absolute bioavailability of intranasal administration was not quantified, and some EVs may enter the periphery via the lymphatic or circulatory systems. Furthermore, as natural EVs containing thousands of bioactive molecules, the therapeutic effects of TH⁺/FOXA2⁺ cell-derived EVs are likely attributable to multi-target-, network-based-synergistic actions rather than a single molecule or pathway. This complexity represents both a potential advantage of EV-based-therapy and a challenge for precise mechanistic dissection. Future studies are warranted to further identify the key effector molecules through cargo knockout or synthetic EV strategies. In summary, this study provides the first evidence, through animal experiments, organoid models, and multi-omics-integration, for the therapeutic potential of TH⁺/FOXA2⁺ cell-derived EVs in the treatment of Parkinson’s disease. The unique molecular cargo carried by TH⁺/FOXA2⁺ cell-derived EVs confers specific therapeutic advantages. Their favorable preliminary safety profile, combined with human-specific-data from the MBO model, greatly enhances the translational value of this study. Future research will focus on elucidating the key therapeutic mechanisms of TH⁺/FOXA2⁺ cell-derived EVs, paving the way for precise and efficient treatment of PD.

## Declarations

### Ethics approval and consent to participate

The animal study protocol was approved by the Animal Ethics Committee of Shanghai Botaiyuekang Life Science Co., Ltd. (Animal Ethics Approval: YK20250901).All animal procedures were performed in accordance with the relevant guidelines and regulations. The human induced pluripotent stem cells (iPSCs) used in this study were purchased from iXcells Biotechnologies USA, Inc. Original somatic cells were collected from healthy donors after obtaining written informed consent. The collection and reprogramming protocols were approved by the relevant institution.

## Consent for publication

Not applicable

## Availability of data and materials

The data will be deposited in the NCBI Gene Expression Omnibus (GEO) repository. These data are available from the corresponding author upon reasonable request prior to public release.

## Competing interests

All authors (including the corporate-affiliated authors) are independent of any commercial interests, and there are no financial or personal relationships that could compromise the authenticity and objectivity of the research findings. All authors take full responsibility for the authenticity and completeness of the research content and data, and declare no other potential conflicts of interest.

## Acknowledgements

Not applicable

## Artificial Intelligence

This manuscript has not used AI-generated content.

## Notes

### Competing Interest Statement

The authors have declared no competing interest.

### Summary of Updates

Changed the order of authors in the system to keep it consistent with that in the text.

